# Joint estimation of neural sources and their functional connections from MEG data

**DOI:** 10.1101/2020.10.04.325563

**Authors:** Narayan Puthanmadam Subramaniyam, Filip Tronarp, Simo Särkkä, Lauri Parkkonen

**Affiliations:** Department of Neuroscience and Biomedical Engineering, Aalto University; Department of Electrical Engineering and Automation, Aalto University

## Abstract

Current techniques to estimate directed functional connectivity from magnetoencephalography (MEG) signals involve two sequential steps; 1) Estimation of the sources and their amplitude time series from the MEG data by solving the inverse problem, and 2) fitting a multivariate autoregressive (MVAR) model to these time series for the estimation of AR coefficients, which reflect the directed interactions between the sources. However, such a sequential approach is not optimal since i) source estimation algorithms typically assume that the sources are independent, ii) the information provided by the connectivity structure is not used to inform the estimation of source amplitudes, and iii) the limited spatial resolution of source estimates often leads to spurious connectivity due to spatial leakage.

Here, we present an algorithm to jointly estimate the source and connectivity parameters using Bayesian filtering, which does not require anatomical constraints in form of structural connectivity or a-priori specified regions-of-interest. By formulating a state-space model for the locations and amplitudes of a given number of sources, we show that estimation of functional connectivity can be reduced to a system identification problem. We derive a solution to this problem using a variant of the expectation–maximization (EM) algorithm known as stochastic approximation EM (SAEM).

Compared to the traditional two-step approach, the joint approach using the SAEM algorithm provides a more accurate reconstruction of connectivity parameters, which we show with a connectivity benchmark simulation as well as with an electrocorticography-based simulation of MEG data. Using real MEG responses to visually presented faces in 16 subjects, we also demonstrate that our method gives source and connectivity estimates that are both physiologically plausible and largely consistent across subjects. In conclusion, the proposed joint-estimation approach based on the SAEM algorithm outperforms the traditional two-step approach in determining functional connectivity structure in MEG data.

## 1 Introduction

The two fundamental and contrasting principles of brain organization are functional segregation and integration, which span multiple spatio–temporal scales. While functional segregation refers to the existence of functionally specialized brain areas that are anatomically separated, functional integration refers to the coordinated interaction between these specialized brain regions, which is essential for various cognitive and perceptual tasks [2]. The investigation of functional relationships between brain regions has been a major task in neuroscience since the beginning of electroencephalography (EEG). It has been hypothesized that neuronal oscillations and their inter-regional synchronization is essential to normal brain function. Being able to measure and characterize brain signals reflecting functional integration shall help us understand how connections between brain regions mediate information and how such connections change e.g., due to learning or to a neurodegenerative disease.

Functional integration is usually described in terms of functional connectivity [9], which is defined as a statistical interdependency of activation dynamics in distinct brain regions. Activation dynamics can be measured indirectly with functional magnetic resonance imaging (fMRI) as fluctuations of the blood-oxygen-level dependent signal or directly with magnetoencephalography (MEG) or EEG as changes in the electromagnetic signals emitted by electrically active neurons. Owing to their millisecond-range temporal resolution, EEG and MEG are ideal methods to monitor the dynamic neural signals.

However, due to their limited spatial resolution, estimating functional connectivity directly from MEG or EEG is not a straightforward task and it typically comprises two sequential steps. In the first step, the ill-posed inverse problem of estimating the source activities from MEG and/or EEG signals is solved, for instance, using minimum-norm estimation (MNE) [20] or beamforming [33, 39]. In the second step, regions-of-interest (ROIs) based on anatomical atlases or separate functional localizer measurements are defined and the estimated activities of all sources in a given ROI are collapsed to obtain representative activity dynamics for each ROI. Functional connectivity can then be estimated between a subset or all the ROIs using a variety of methods. While measures such as coherence, phase-locking value, phase-lag index – to mention a few – yield a symmetric connectivity matrix, approaches based on multivariate autoregressive (MVAR) models can result in a directed or asymmetric connectivity matrix; partial directed coherence (PDC) and directed transfer function are such measures. With these directed measures, the sources (or ROIs) are typically assumed to be few and their locations known.

A major weakness of the two-step approach is that, since the estimation of source activities and the connectivity between them occur in two distinct steps, the connectivity estimation method is not informed about the assumptions the source estimation algorithm makes, e.g. independence across the sources, which eventually leads to biased connectivity estimates. In particular, when distributed source estimation algorithms – such as MNE – are used, two-step approaches often lead to spurious connections between sources.

To include additional information in source estimation, linear MVAR models with spatially local interactions and self-interactions have been proposed [11, 13, 24]. However, such approaches ignore long-range interactions across brain regions, which are the hallmark of functional integration.

Joint estimation of source activities and connectivity has been popularly addressed by a confirmatory approach known as dynamic causal modeling (DCM) [10]. In DCM, ROIs are pre-specified and the activity in each ROI is modeled through a set of nonlinear differential equations. Then, the estimated connectivity solutions for plausible generative models are compared using Bayesian model selection and an appropriate model is chosen.

Over the last few years, data-driven, exploratory approach has been used for addressing the joint estimation problem using MVAR models [4, 35]. In this approach, expectation–maximization (EM) approach with a Kalman smoother in a state-space formulation is used to obtain a maximum a-posteriori (MAP) estimate of the source signals and a maximum likelihood (ML) estimate of the MVAR coefficients, which describe the interaction. However, these approaches still require the ROIs or source locations to be predefined. Recently, Fukushima and colleagues [12] proposed joint estimation of whole-brain connectivity from MEG signals without requiring ROIs to be predefined but relying on structural connectivity information, for example, from diffusion tensor imaging (DTI). However, such information may not be available, and functional connectivity patterns typically only loosely follow structural connectivity.

In this work, we propose a framework for the joint estimation of source locations and their directed functional connections from MEG data. We formulate a state-space model (SSM) for the evolution of source locations and as well as of source amplitudes. The transition model for source locations is a first-order random walk, while for source amplitudes it is a *P*−th order MVAR model, with the coefficients representing the interaction between the sources. The MEG measurement depends nonlinearly on the source location but linearly on the source amplitude, which are embedded in our measurement model. Estimation of the MVAR coefficient matrix is then reduced to a system identification problem in a non-linear state space with a tractable linear sub-structure, which can be solved efficiently using stochastic approximation expectation–maximization (SAEM) approach combined with Rao–Blackwellization [36]. Once the MVAR matrix is estimated, directed functional connectivity estimates can then be obtained using methods such as generalized partial directed coherence (gPDC) [1] which provides a frequency-domain representation of the MVAR model.

To evaluate the performance of our method – referred to as SAEM from now on – against two-step approaches, we used MVAR-based as well as electrocorticography (ECoG) -based simulations, where we also incorporated modelling errors. The MVAR simulation is based on the Berlin brain connectivity benchmark (BBCB) by Haufe and colleagues [16] and it comprises several scenarios with two sources that are either non-interacting (Type-0) or that have a linear, unidirectional interaction (Type-I); see Section 2.4 for details. Our ECoG-inspired simulations are based on an open dataset that contains responses to various visual stimuli; we projected ECoG signals at electrodes in the right superior occipital and inferior occipital areas to MEG sensors according to the MEG forward model (see Section 2.5 for details). As the real MEG data, we used publicly available recordings [40], where the subjects (*N* = 16) were shown human faces.

For source estimation in the two-step approach, we used a distributed source estimation method called exact low-resolution brain electromagnetic tomography (eLORETA) and two spatial filtering methods: linearly constrained minimum variance (LCMV) and multiple constrained minimum variance (MCMV) beamformers. In the second step, we fitted an MVAR model to estimate the interaction coefficients and transformed them in to a gPDC matrix that gives directed functional connectivity. For the two-step approaches, we assumed the source locations to be known, except for the MCMV method against which we also evaluated the source-localization performance of our approach using the ECoG-based simulations.

## 2 Materials and methods

### 2.1 Notation and assumptions

Throughout this paper, vectors or matrices are referred to in bold fonts. The symbols *𝔼*[·] and Cov[·] are used to represent the expectation and covariance operators respectively. *p*(**x**) is the probability density function (pdf) of **x** and *p*(**x**|**y**) for the pdf of **x** given **y**. *𝒩* (**x**|***µ*, Σ**) represents a multivariate Gaussian distribution of **x** with mean ***µ*** and covariance **Σ. *µ***_*t*|1:*t*−1_ and ***µ***_*t*|1:*t*_ refer to the predicted and filtered mean, respectively, whereas ***µ***_*t*|1:*T*_ refers to the smoothed mean of the Kalman filter. In a similar vein, we define **P**_*t*|1:*t*−1_, **P**_*t*|1:*t*_ and **P**_*t*|1:*T*_ for the Kalman filter covariance. The symbol ∼ is used to denote drawn from a distribution. For any sequence 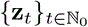, we write **z**_*m*:*n*_ ≜ {**z**_*m*_ … **z**_*n*_}, where *m > n* and *m, n ∈* ℕ_0_.

We denote the number of MEG time samples as *T* and number of trials as *J*. The *M*-channel MEG measurement of the *j*-th trial at time *t* is denoted as 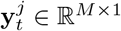. We assume that only a reasonable number of sources (1 < *N*_*s*_ ≤ 5) are active and they can be modeled as dipoles oriented normal to the local cortical surface. We further assume that the source locations remain constant across the trials. We denote the location of *N*_*s*_ sources at time *t* as 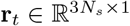. The amplitudes of *N*_*s*_ sources at time *t* and trial *j* are denoted as 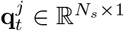. We define **x**_1:*t*_ ≜ {**r**_1:*t*_, **q**_1:*t*_}. Note that 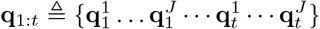. In a similar vein, we define **y**_1:*t*_. Furthermore, we assume that the number sources is known a priori, and all the noise in the model has a Gaussian distribution with zero mean.

### 2.2 State-space model

We formulate a SSM of the form

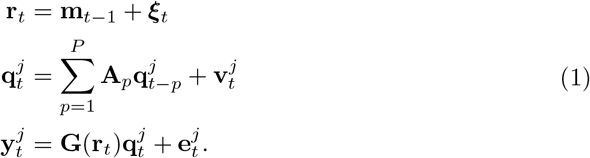

The matrix **A**_*p*_ consists of the AR coefficients representing the correlation between the sources at time-delay *t* − *p*, across all the trials. 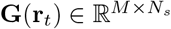 is the lead-field matrix describing the contribution of *N*_*s*_ dipoles of unit strength at location **r**_*t*_ to *M* MEG sensors. The process noise for the source amplitudes is given by 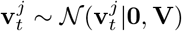. The measurement noise is denoted by 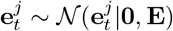.

Note that the source locations do not depend on individual trials but are rather estimated from the average across the trials (see Section 2.3.2). We incorporate an artificial evolution for source locations in our SSM, since we use a particle filter to estimate them. However, as it is commonly assumed in several dipole-localization approaches, the true locations of the sources are fixed in time and only their amplitudes vary. To estimate fixed source locations using a particle filter, we adopt a kernel smoothing approach via shrinkage described in Ref. [27]. The kernel location is given by **m**_*t*−1_ and is specified using a shrinkage rule. The artificial evolution noise for source locations is given as ***ξ***_*t*_ ∼*𝒩* (***ξ***_*t*_|**0**, *h*^2^**Ξ**), where *h* is the smoothing parameter. The kernel smoothing approach is further explained in the supplementary section.

To facilitate the application of Bayesian filtering algorithms, a *P*−th order MVAR model has to be reformulated into a first-order MVAR model (see SI) as follows:

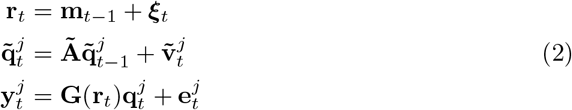

with

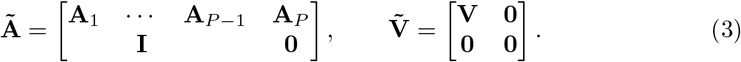

### 2.3 Stochastic approximation Expectation–Maximization

Given a latent variable **x**_1:*T*_ and measurement **y**_1:*T*_, an EM algorithm employs an iterative scheme where the auxiliary quantity

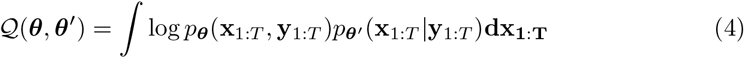

related to the lower bound of the marginal log-likelihood is maximized as opposed to the direct maximization of the marginal log-likelihood, which is not always possible. Starting from an initial guess ***θ***[0] *∈* **Θ**, an EM algorithm iterates between the two steps until convergence or the maximum number of iterations is reached:

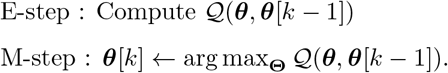

At every *k*-th iteration of the EM algorithm, we also get the full posterior or the joint smoothing distribution (JSD), *p*_***θ***[*k*−1]_(**x**_1:*T*_|**y**_1:*T*_) which solves the state inference problem related to the estimation of source locations and strength. In the case of nonlinear state-space models, the E-step does not have a closed-form solution and needs to be approximated using a stochastic version of the EM algorithm, where Monte-Carlo approximation based on simulations can be used to approximate the the integral in Equation 4 [23]. However, such methods make inefficient use of the simulations, as at each new iteration the values simulated during the previous iteration are completely discarded.

Deylon and colleagues [6] proposed an alternative scheme, known as the stochastic approximation EM (SAEM) algorithm, where the simulated variables are gradually discarded with each iteration using a forgetting factor *ζ*_*k*_. In SAEM, the E-step is replaced by a stochastic E-step:

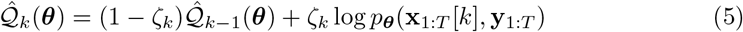

where ***x***_*1:T*_ [*k*] ∼ *p*_***θ***[*k*−1]_(**x**_1:*T*_ |**y**_1:*T*_) and {*ζ*_*k*_}_*k≥*1_ is a decreasing sequence of positive step-size satisfying the constraints ∑_*k*_ *ζ*_*k*_ = *∞* and 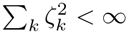 [25].

#### 2.3.1 Rao–Blackwellization

Given the MEG observations **y**_1:*T*_ and the number of sources *N*_*s*_, we are interested in estimating the state **x**_1:*T*_ comprising the location **r**_1:*T*_ and amplitude 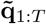 of the sources, and the parameters, AR coefficient matrix **Ã**, process noise covariance matrix for the source amplitudes 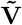 and measurement noise covariance matrix **E**. We can denote the parameters to be estimated as 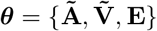.

Making use of conditional probabilities, we can express the target distribution of interest at SAEM iteration *k* as

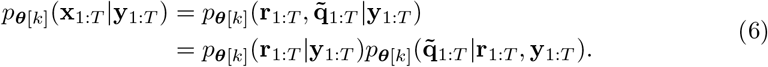

Since the distribution 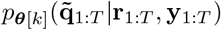 is Gaussian, the inference of the conditionally linear states 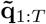 can be done in closed form using a Kalman filter and smoother [32], thus requiring Monte-Carlo-based simulation method only to approximate *p*_***θ***[*k*]_(**r**_1:*T*_|**y**_1:*T*_). Such marginalization reduces the asymptotic variance of the estimator and is known as Rao–Blackwellization [3, 32].

As mentioned previously, we will use the trial-averaged MEG data

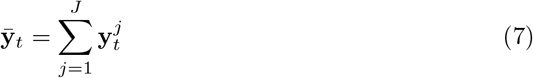

to estimate the source location. Thus, the posterior distribution for source locations is given as 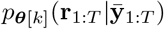. In particular, we will use particle Markov-Chain Monte-Carlo method to approximate the distribution 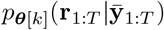 infer the source location, and a Kalman smoother to estimate the posterior 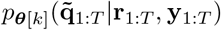 is used to infer the amplitude. We refer to this approach of inferring the source locations and amplitudes as Rao–Blackwellized PMCMC (RB-PMCMC) approach [25].

#### 2.3.2 Estimating source locations

For our model, direct sampling of **r**_1:*T*_ [*k*] from 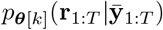 is not possible. We resort to a MCMC technique and propose a Markov kernel Π_***θ***[*k*−1]_(·), such that it leaves the posterior distribution 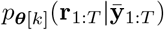 invariant. Thus, instead of sampling directly from the posterior distribution 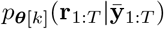, it is sufficient to generate samples using a Markov kernel, which is assumed to be ergodic with stationary distribution 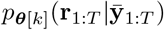. Let Π_***θ***_(·) be such a kernel. We then have

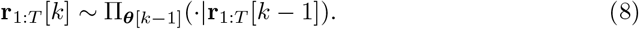

By simulating a Markov chain with the kernel Π_***θ***_(·), the marginal distribution of the chain will approach the posterior distribution 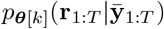 for sufficiently large *k* [36]. PMCMC approaches can be used to construct an efficient and high-dimensional kernel Π_***θ***_(·). In PMCMC, a sequential Monte Carlo (SMC) sampler such as a particle filter is used as the Markov kernel. As proposed by Lindsten and colleagues [25], we use a variant of the particle filter known as conditional particle filter with ancestor sampling (CPF-AS) as the kernel. Using CPF-AS, we approximate the posterior distribution at iteration 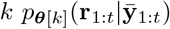 as [26]

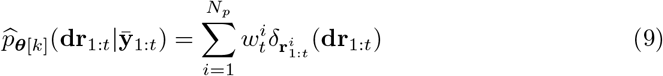

where *δ*(·) is an impulse function and 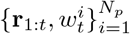 is a weighted particle system with *N*_*p*_ particles such that 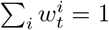. Details regarding the CPF-AS method is given in the supplementary text.

#### 2.3.3 Estimating source amplitudes

Conditioned on the source locations, 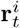 for *i* = 1, 2, …, *N*_*p*_, the predictive distribution of 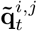 is given as

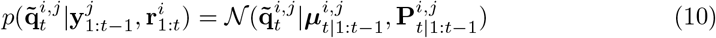

and the filtering distribution of 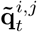 is given as

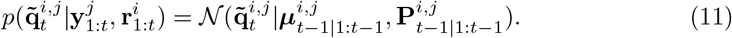

We use a Rauch–Tung–Striebel (RTS) smoother to obtain the marginal smoothing distribution of 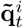, which is given as

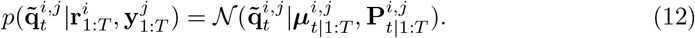

From the RTS smoother, we also obtain the one-lag Kalman smoother covariance, 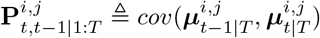. See supplementary text for the explicit expressions of the mean and covariance of the Kalman filter and smoother.

At iteration *k* of the SAEM algorithm, we update 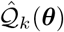 in the stochastic E-step given in Equation 5 by making use of all the particles given by CPF-AS instead of just using **r**_1:*T*_ [*k* + 1] [25]. That is,

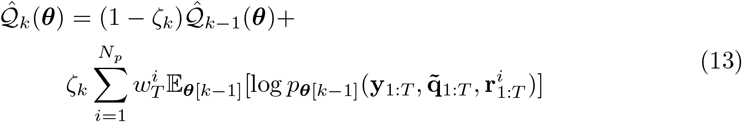

where the 𝔼_***θ***[*k*−1]_[·] is w.r.t the linear state 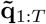, which is obtained by the Kalman smoother. For notational convenience, we refer to 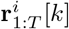 and 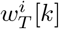 as simply 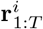 and 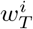.

#### 2.3.4 ML estimation of parameters

Connectivity and noise covariance parameter estimates are obtained in the maximization step of the SAEM method, where our aim is to maximize 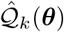 given in Equation 13, with respect to 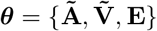. The complete log-likelihood log 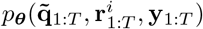 in Equation 13 can be factorized as

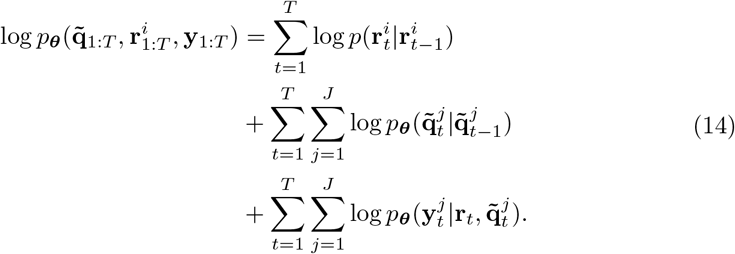

Only the second and third term in the above expression depend on ***θ*** and thus taking the expectation of only this part of the joint log-likelihood, we have [32]:

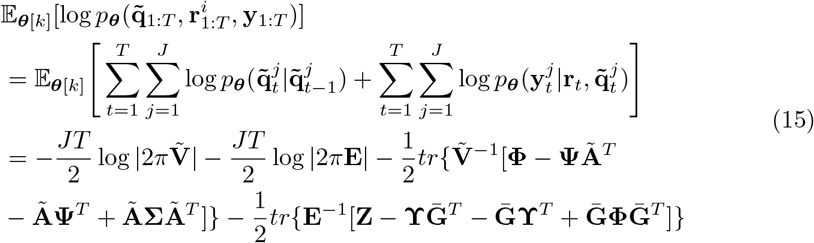

where 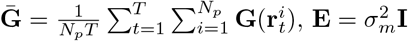 and

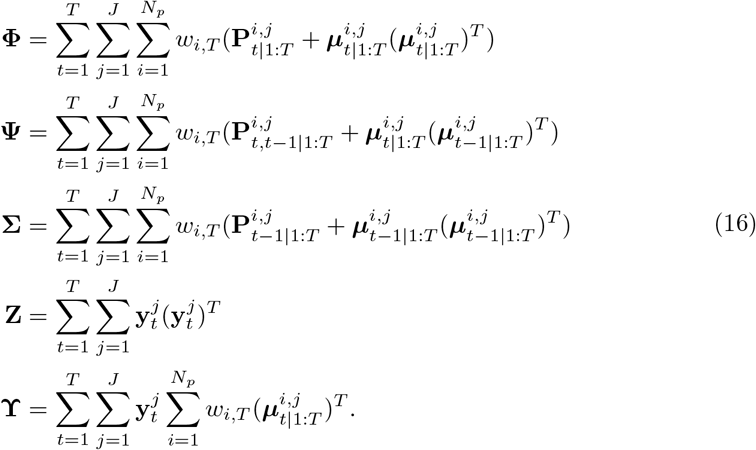

Collecting the sufficient statistics computed above using the Kalman smoother into a matrix

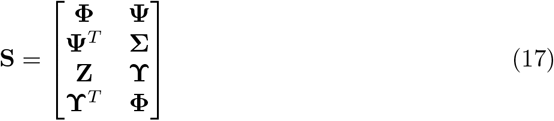

the computation of the stochastic E-step given in Equation 13 reduces to recursively updating

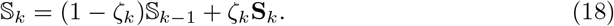

At iteration *k*, maximizing 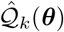 w.r.t **Ã**, 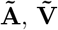and **E** [32] (see supplementary text for details) leads to the following update equations:

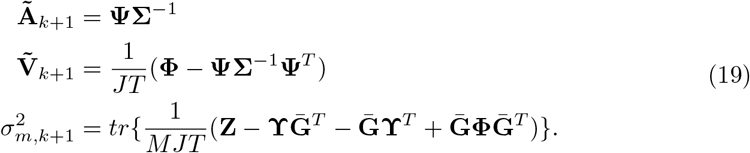

We also account for the brain noise in our SSM by modifying the measurement equation to

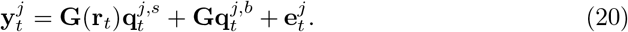

The noise covariance is now given as 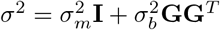, where the additional term 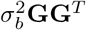 represents the brain noise projected to the MEG sensors. Note that we assume brain noise to be white, i.e, 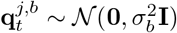. In this scenario, the closed-form update for 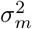 and 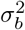 no longer exists and we use gradient-descent algorithm with backtracking to update 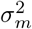 and 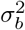. The algorithm for the SAEM method and the RB-PMCMC method within it is given in the supplementary text.

Since the SAEM algorithm finds only a local optimum, we use multiple initializations (20 for the ECoG-based simulations and real data) for the SAEM algorithm and select the solution that that gives the maximum log-likelihood (see Equation 26).

### 2.4 MVAR Simulations

We designed our first simulations according to the Berlin brain connectivity benchmark (BBCB) proposed by Haufe and colleagues [16]. The BBCB benchmark is based on a head model originally consisting of 75000 vertices on the cortical surface, divided into eight ROIs. For the purpose of the simulation and reasonable computational cost, the BBCB framework uses the downsampled version with 2000 vertices. Two sources, modeled as current dipoles oriented normal to the local cortical surface, were placed randomly in the source space, with the only constraint that they should be at least 3 cm apart. Two scenarios were considered 1) Type-0: No interaction between the sources, and 2) Type-I: Unidirectional interaction between the sources. We used a second-order MVAR model to simulate these scenarios. Brain noise was added by randomly selecting 500 source locations and modelling their activity as uncorrelated pink noise, which was then projected to the sensors and added to the brain signal produced by the two dipoles. In addition to the brain noise, we also added Gaussian measurement noise. Thus, the simulated MEG signal (*T* = 300 samples, *J* = 1 trial) is given as

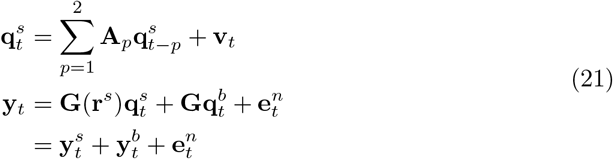

where 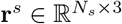 are the locations of the *N*_*s*_ = 2 dipoles generating brain signal and 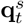 are the corresponding amplitudes. The measurement noise is generated according 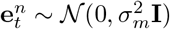.

We fixed SNR_meas_ at 5 and added brain noise at SNR 1, 3, 5 10, where brain noise SNR is defined as

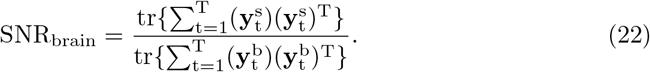

Finally, the SNR_meas_ in this scenario becomes

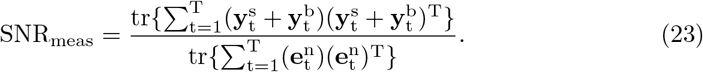

We generated 50 realizations for each value of SNR_brain_ and each connectivity scenario (non-interacting and interacting sources).

#### 2.4.1 Performance evaluation

We evaluated the performance of our proposed SAEM algorithm on three criteria - 1) Source localization error (SLE), 2) Relative error (RE) for the gPDC matrix and noise covariance, and 3) Estimation of the direction of interaction. We also compared the performance of our proposed method with standard two step-approach of using distributed source imaging method eLORETA and spatial filtering methods LCMV and MCMV beamformer followed by AR model fitting using Vierra–Morf (VM) algorithm.

For the SAEM approach approach, the source location is given by the output particle MCMC smoother (see Section 2.3.2), which we average over the number of particles and time points. We consider source localization to be successful if the localization error is less than 2 cm between the estimated source location and the true true location. Only in the case that both sources are correctly localized according to the aforementioned definition, the estimation of AR and covariance matrices as given by the maximization step of the SAEM algorithm is considered.

For the two-step approach, we assume that the source location is known a-priori and thus only estimate the source amplitudes and the AR matrix using the VM algorithm. Finally, in order to estimate the direction of interaction and consequently compute the true positive rate (TPR) and false positive rate (FPR), we compute the gPDC. We used causal Fourier transform (CFT) surrogates (100 surrogates, significance level 0.01) to find statistically significant gPDC values [7].

We define the number of true positives (#TP) as the number of correctly identified links and the number of false negatives (#FN) as the number of missed links. The number of false positives (#FP) is defined as the number of incorrectly estimated links and the number of true negatives (#TN) as the number of correctly identified non-links. Thus, for the unidirectional interaction case, we defined both TPR and FPR, while for the non-interacting scenario, only FPR was defined. Thus, TPR and FPR are given as

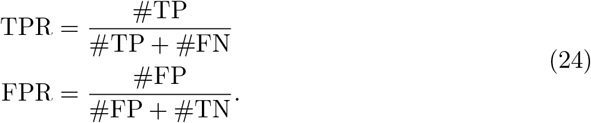

Note that in case of the SAEM method, TPR and FPR were only computed for those cases where both sources were correctly identified. We used bootstrapping technique to obtain the mean and standard deviation (SD) for the performance metrics SLE, TPR, FPR, gPDC RE and *σ*_*m*_ RE as follows: We generated *B* = 100 sets of bootstrap samples of size *S* = 50, obtained by sampling with replacement from the original 50 simulations. In case of SLE, we first computed the average localization error, 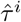, over the two sources for each simulation *i* within the bootstrap sample and then computed the mean over the bootstrap sample 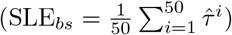. Finally, the mean SLE over *B* bootstrap sets was computed as 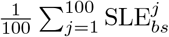. In case of other performance metrics, we simply computed the mean value of the metric; first we evaluated each bootstrap sample and then computed the mean of these mean values (obtained for each bootstrap sample) over the *B* bootstrap sets. The SD was computed in an analogous manner.

### 2.5 Simulation using electrocorticography signals

Although MVAR simulations are useful as the ground truth is known, they do not capture the nonlinear characteristics of neural signals. We used electrocorticographic (ECoG) recordings (available at http://www.fieldtriptoolbox.org/example/ecog_ny/) as the basis for a realistic simulation of MEG signals for which we still have information on the ground truth. Specifically, the employed dataset was from a paradigm where the subject was shown images of false fonts, houses, other objects, textures, bodies, text or faces. For this study, we selected only the trials pertaining to the face stimuli. The ECoG grid had 128 electrodes in total, out of which we selected two electrodes located in the right occipital and the right inferior inferior temporal areas where the electrodes were most responsive to the face stimuli (electrodes SO3 and IO2 in the original data. The locations of these two electrodes were approximately mapped to source space determined for the head models of the 16 subjects and wer place in the right lateral occipital parcel (source corresponding to the electrode SO3) and right fusiform parcel (source corresponding to the electrode IO2) of the Desikan–Killiany (DK) atlas. The ECoG signals were projected to the MEG sensors using the MEG forward model (see Section 2.6.2). To account for modelling errors, we used a single-layer boundary element model (BEM) model with icosahedron subdivision to 5120 nodes (using “ico4” in MNE-Python; [14]) for data generation and for estimation we used a BEM model with 1280 nodes (“ico3” in MNE-Python). Measurement and brain noise was added at SNR_meas_ = 3 and SNR_brain_ = 1, respectively, to the simulated MEG signals (see section 2.4). Brain noise was modelled as uncorrelated pink noise generated from randomly-selected 2000 vertices of the source space and projected to the MEG sensors, while the measurement noise was modelled as a Gaussian. Furthermore, to make the simulation scenario more realistic, we set the number of sources, *N*_*s*_, to be estimated by the SAEM and MCMV algorithms to 5. In reality, we usually do not know the number of sources and thus setting it to a reasonable number is in line with our assumptions (see Section 2.1). For the MCMV-based source localization, we used the multi-source activity index (MAI) with identity covariance matrix [29]. The gPDC computed after fitting an MVAR model to the ECoG signal from the two pre-selected electrodes was considered as the ground truth for the connectivity between the two sources.

### 2.6 Real MEG data

We used the data made publicly available by Wakeman and Henson [40] (available at https://legacy.openfmri.org/dataset/ds000117/) and consisting of simultaneous MEG (Elekta Neuromag Vectorview, 204 gradiometers and 102 magnetometers; MEGIN Oy, Helsinki, Finland) and EEG recordings (70 channels) from 19 healthy subjects when viewing famous, non-famous and scrambled faces. For details on data acquisition and experimental design, we direct the reader to the original publication. Briefly, the subjects were presented with 300 grey-scale photographs; 150 faces of famous people and 150 faces of non-famous (unknown to the subjects) people. In addition, 150 images of scrambled famous or unknown faces were also presented. Each image was presented twice (either immediately or after 5–15 intervening images). For the sake of simplicity in our analysis, we did not distinguish between the initial and repeated presentations and thus there are three trial types: famous, non-famous and scrambled faces. For the purpose of this study, we used only the MEG magnetometer recordings (102 channels) and considered only the trials with non-famous and scrambled faces presented. Signal Space Separation (SSS) [37] was already applied to the MEG data for the suppression of magnetic interference.

#### 2.6.1 Preprocessing

Due to issues with the data quality, three subjects were excluded (Subjects 1, 5 and 19 in the original dataset). The MEG data of the remaining 16 subjects were preprocessed using MNE-Python [14], following the guidelines published by Jas and colleagues [22]. Briefly, the data were band-pass-filtered (zero-phase finite impulse response filter) to 0.3–40 Hz and down-sampled to 220 Hz. Artifacts such as eye-blinks and heart beats were removed using independent component analysis (ICA), with the maximum number of components to be removed set to 2 for the ocular artifact and 3 for the cardiac artifact. Epochs were extracted –200 … 2900 ms with respect to the stimulus onset, and the baseline period was also used to estimate the measurement noise covariance using the shrinkage method. Rejection threshold for the trials were set to 4 × 10^−12^ fT for magnetometers and 4 × 10^−10^ fT/cm for gradiometers. We chose 100 trials randomly for the unfamiliar face stimuli and considered the data between 0 and 250 ms for the SAEM algorithm to estimate source and connectivity parameters.

#### 2.6.2 Head modelling

We segmented the MRI for the brain volume and for the cortical mantle using the FreeSurfer software (version 5.1.0) [31] with the ‘recon-all’ command. We defined the source space by octahedron subdivision (“oct6” in MNE-Python), resulting in about 4098 vertices per hemisphere. For each subject, we used a single-layer boundary element method (BEM) model comprising of just the inner skull to compute the forward solution. We excluded source points that were less than 5 mm to the inner-skull surface. Using the forward solution, the lead-field was obtained for each MEG sensor, assuming source orientation normal to the local cortical surface.

##### Model order and the number of sources

To estimate the model order, we used Bayesian information criteria (BIC) and varied *P* from 1 to 20. In BIC, the aim is to find the *P* at which the following function attains its minimum:

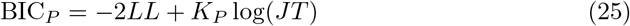

where the log-likelihood, *LL*, is given as

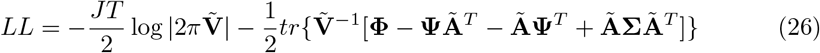

and 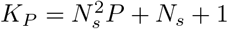 is the number of free parameters. Here 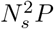 refers to the number of free parameters in **Ã**, *N*_*s*_ refers to the number of free parameters in 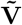 (i.e., only the diagonal entries of **V**) and one free parameter for the brain noise variance 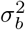. We estimated the measurement noise covariance from the baseline data (−300 to 0 ms). We also whitened the data and as a consequence 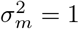.

Using BIC, we found that the model order varied between 8 and 13 across subjects. We chose model order 10 as for most subjects the chance in BIC after *P* = 10 was negligible. We set the number of sources for each subject to 5. This is a reasonable assumption as it is known that the sources underlying the processing of face stimuli are located in the occipital, temporal and probably frontal cortices, which refer to the core and the extended system [17].

## 3 Results

### 3.1 MVAR-based simulations

Below we summarize the results from applying our novel SAEM algorithm as well as the reference two-step approaches to simulations comprising non-interacting and interacting sources at several signal-to-noise ratios.

#### 3.1.1 Source localization

Table 1 shows the mean source localization error (SLE) for the SAEM method applied to the simulations with non-interacting (Type-0) and unidirectionally interacting (Type-I) sources. The mean SLE is under 20 mm at all simulated levels of SNR_brain_ for both Type-0 and Type-I scenarios, with the mean SLE decreasing as SNR_brain_ increases.

**Table 1.** SAEM algorithm applied to MVAR simulations. Mean source localization error (SLE) and the percentage of sources localized with SLE ¡ 20 mm (CH) across a range of brain-signal-to-noise ratios. The values are given for non-interacting (Type-0) and unidirectionally interacting (Type-I) sources. The measurement SNR was fixed at 5. The values are followed by ± standard deviation.

| SNR<sub>brain</sub> | SLE for Type-0<br>mm | CH for Type-0<br>% | SLE for Type-I<br>mm | CH for Type-I<br>% |
| --- | --- | --- | --- | --- |
| 1 | 16.1 ± 2.0 | 43.4 ± 6.2 | 15.9 ± 2.3 | 54.3 ± 7.1 |
| 3 | 5.6 ± 1.8 | 86.3 ± 4.8 | 9.7 ± 2.3 | 81.1 ± 5.5 |
| 5 | 3.2 ± 1.3 | 92.1 ± 3.7 | 6.3 ± 1.5 | 83.7 ± 5.2 |
| 10 | 2.4 ± 1.0 | 94.5 ± 3.5 | 5.1 ± 1.6 | 89.5 ± 4.8 |

Table 1 also shows the mean percentage of cases for which the maximum SLE across two sources was less than 20 mm, which we refer to as the number of correct hits (CH). For both Type-0 and Type-I scenarios, CH is greater than 80% for all values of SNR_brain_ *>* 1 and CHs increase with SNR_brain_.

#### 3.1.2 Connectivity estimation

Figure 1 shows the mean reconstruction error (RE) for the gPDC matrices obtained using the two-step approach (LCMV, MCMV and eLORETA source estimation followed by AR-fitting) and the joint estimation with the SAEM method. When the sources are not interacting (Type-0), the LCMV method followed by MVAR-fitting gives the lowest RE, however, the SAEM method also gives a low RE, which remains < 2% for all values of SNR_brain_. In the case of interacting sources (Type-I), we can see that the SAEM method gives the lowest RE compared to all tested two-step approaches (*p* < 0.001) despite the advantage of the two-step approaches that the true source locations were given and only the source amplitudes and the MVAR matrix had to be estimated.

**Figure 1.**
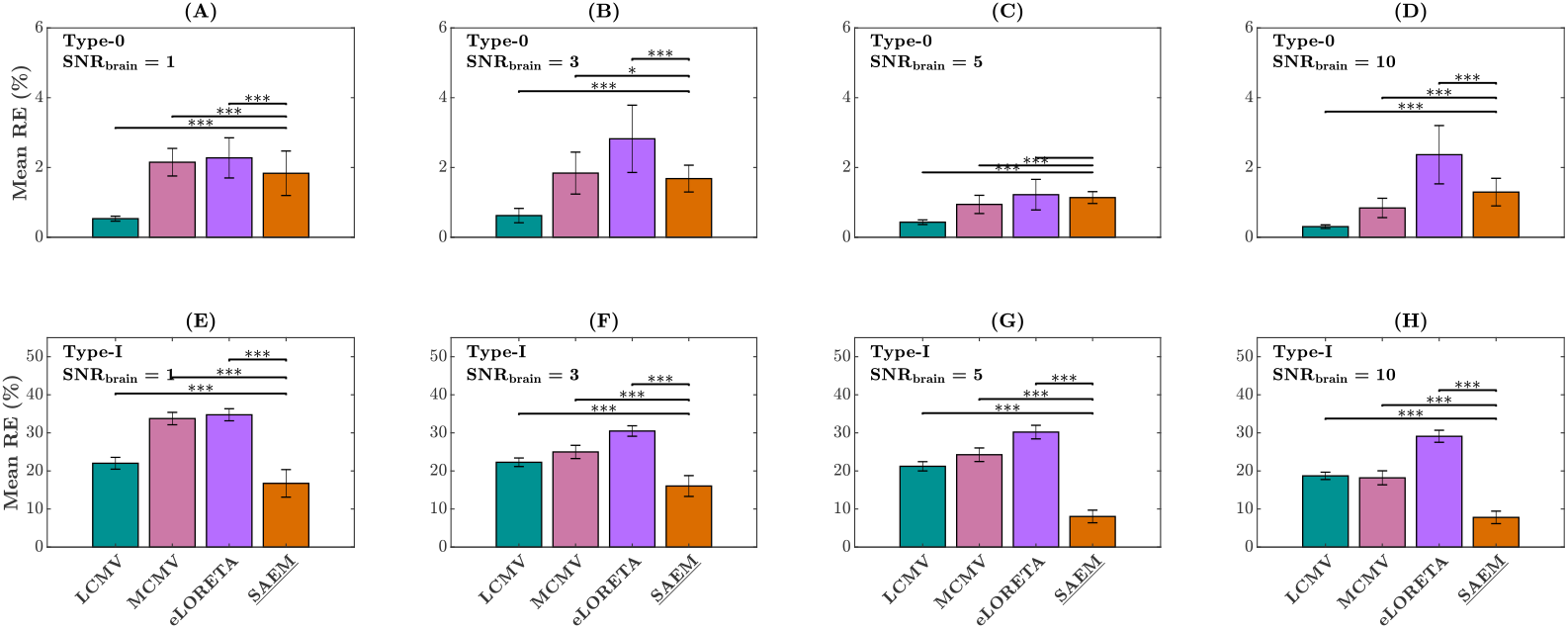
MVAR simulations. Mean reconstruction error of connectivity coefficients at SNR_brain_ = 1, 3, 5 and 10 for non-interacting (A–D) and unidirectionally interacting (E–H) sources. The measurement noise SNR was fixed to 5, and the error bars represent the standard deviation. ^*^*p* < 0.05, ^**^*p* < 0.01 and ^***^*p* < 0.001.

We also computed the mean reconstruction error for the measurement noise (*σ*_*m*_) for different levels of SNR_brain_ and for simulations of Type-0 and Type-I (see Supplementary figure S7). We found that the SAEM algorithm was able to reliably estimate the noise covariance even at low SNR_brain_, with RE of 0.7 ± 0.1% at SNR_brain_ = 1 for both Type-0 and Type-I scenarios.

#### 3.1.3 Direction of interaction

The false positive rates (FPR) of directed connections identified with the two-step approaches (LCMV, MCMV and eLORETA followed by MVAR-fitting) and with the joint estimation approach using the SAEM method are shown in Figure 2 for different levels of SNR_brain_. For the Type-0 scenario, both the LCMV and SAEM method gave a lower FPR (< 15%) than MCMV and eLORETA (*>* 20%). For the Type-I scenario, the joint estimation approach using SAEM gave the lowest FPR compared to all tested two-step approaches for all values of SNR_brain_. Among the two-step approaches, MVAR-fitting of source amplitudes obtained with MCMV outperformed both LCMV and eLORETA. The LCMV approach gave the highest (worst) mean FPR.

**Figure 2.**
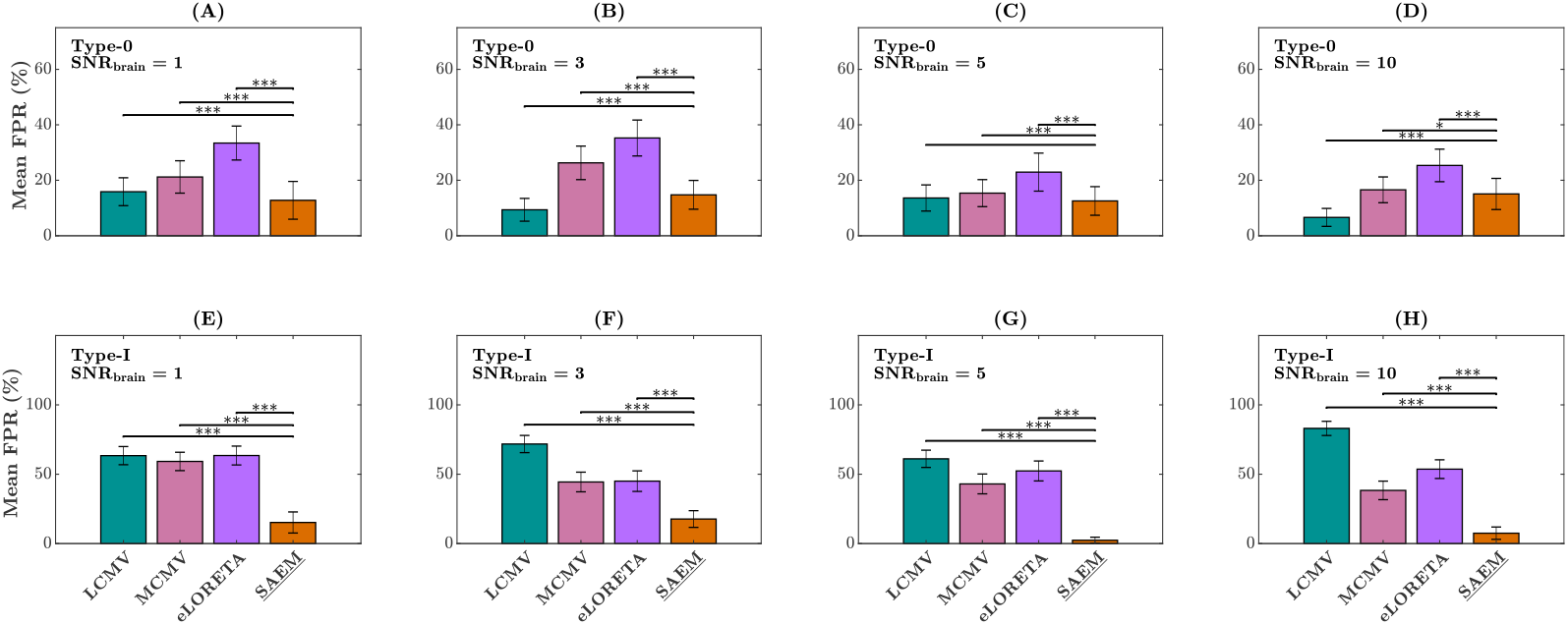
MVAR simulations. Mean false positive rates of identified functional connections at SNR_brain_ = 1, 3, 5 and 10 for non-interacting (A–D) and unidirectionally interacting (E–H) sources. The measurement noise SNR was fixed to 5, and the error bars represent the standard deviation. ^*^*p* < 0.05, ^**^*p* < 0.01 and ^***^*p* < 0.001.

Figure 3(A–D) shows the mean true positive rates (TPR) of identified connections for Type-I scenarios at several SNR_brain_. Although there were statistically significant differences between the methods, joint estimation and all two-step approaches gave rather similar mean TPR for all values of SNR_brain_ when comparing to the substantial differences in FPR results.

**Figure 3.**
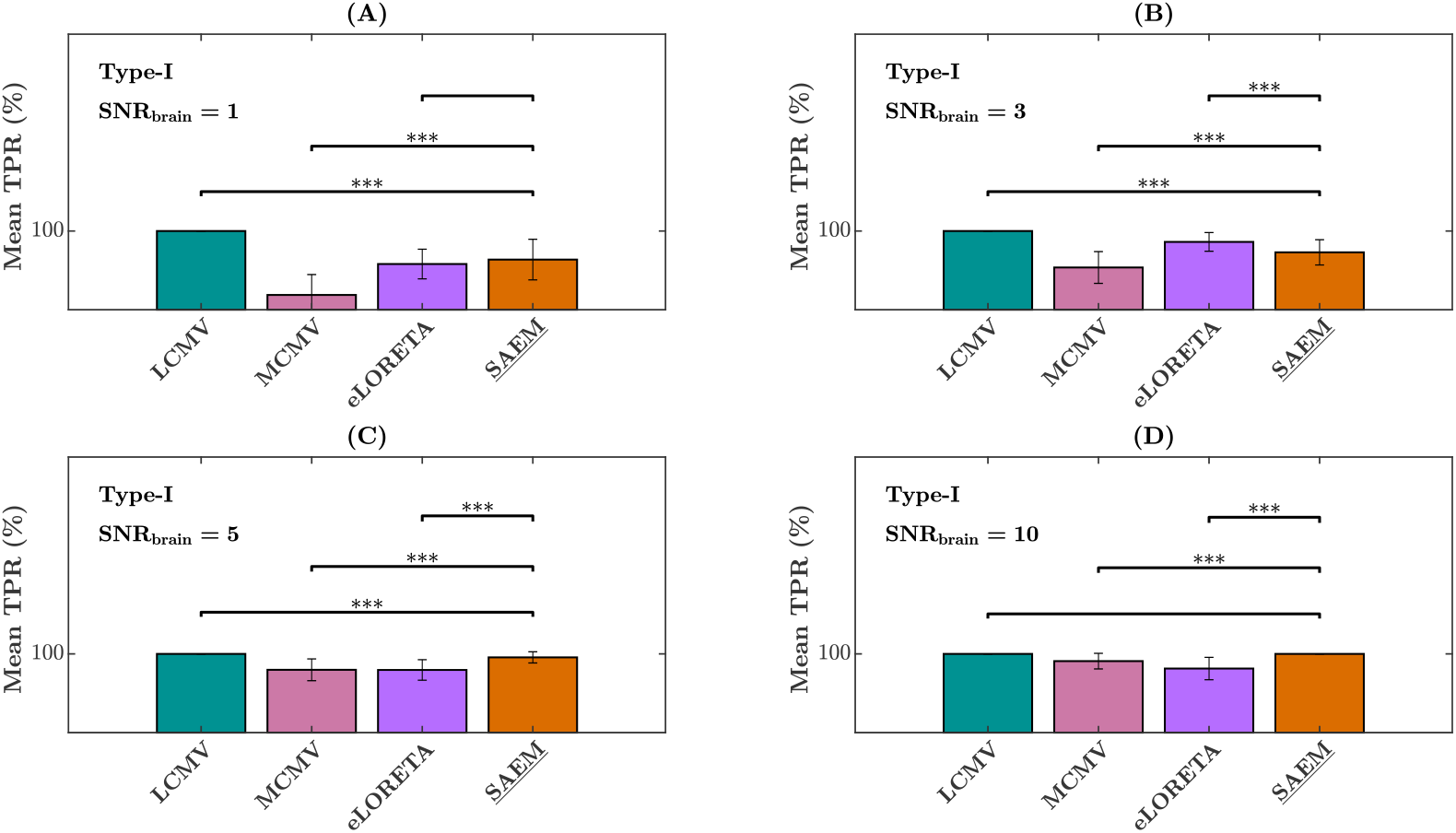
MVAR simulations. Mean true positive rates of identified functional connections at SNR_brain_ = 1, 3, 5 and 10 (A–D) for interacting sources.The measurement noise SNR was fixed to 5, and the error bars represent the standard deviation. ^*^*p* < 0.05, ^**^*p* < 0.01 and ^***^*p* < 0.001.

Further results (estimates of source locations and amplitudes, convergence of parameter estimates and log-likelihood estimation) from applying the SAEM algorithm on MVAR simulations are presented in Figures S1–S6 in the supplementary material.

### 3.2 ECoG-based simulations

We applied the MCMV-based two-step method and our SAEM joint-estimation method to the MEG data simulated based on real ECoG-recorded responses to face stimuli. We estimated locations, amplitudes and connections of the sources jointly with the SAEM method (Fig. 4) and in two consecutive steps using the MCMV method for source localization followed by MVAR fitting of connectivity parameters (Fig. 5). The number of sources was assumed to be 5 for both the SAEM and MCMV methods. We first show qualitative comparisons of the results obtained with the two algorithms and then present a quantitative assessment.

**Figure 4.**
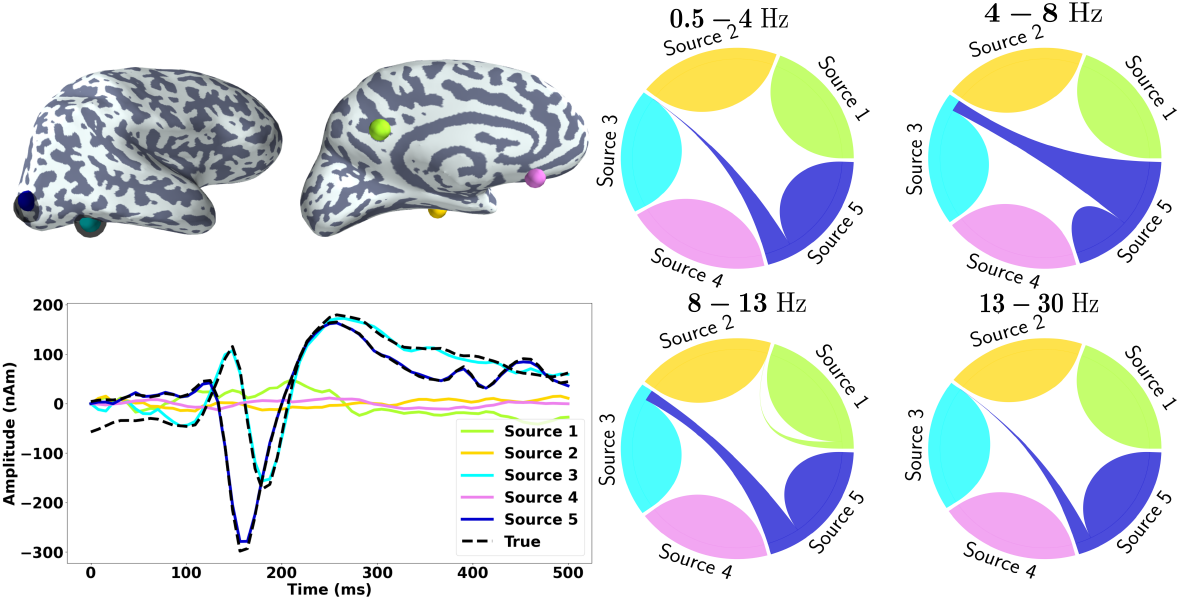
ECoG-based MEG simulations. Estimates of source locations, amplitudes and functional connectivity using the SAEM method in one subject. The true locations (ECoG electrodes) of the two sources are shown as black spheres. The chord plot describes directed functional connectivity between the sources; a shrinking arc from Source A to Source B indicates that Source A is driving Source B.

**Figure 5.**
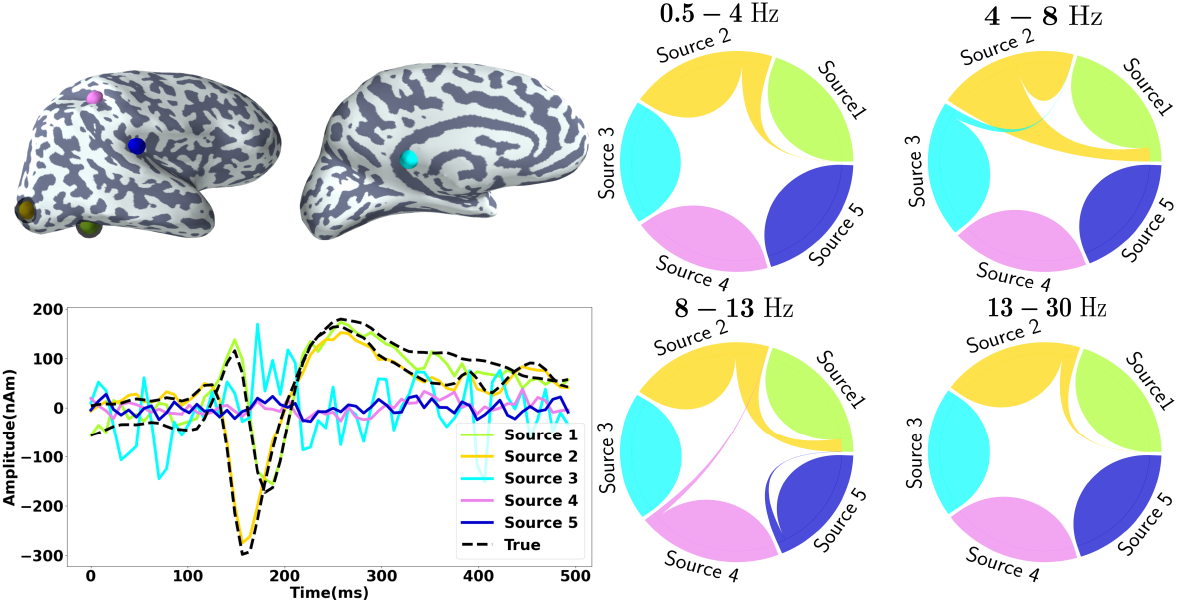
ECoG-based MEG simulations. Estimates of source locations, amplitudes and functional connectivity using MCMV for source localization followed by MVAR fitting in one subject. The true locations of the two sources are shown as black spheres. The chord plot describes directed functional connectivity between the sources; a shrinking arc from Source A to Source B indicates that Source A is driving Source B.

Figure 4 shows that the time-varying amplitudes of two sources (Sources 3 and 5) estimated by the SAEM method match well with the ground truth, i.e., the ECoG measurements. The estimated amplitudes of the three extra sources (Sources 1, 2 and 4) remain close to zero, also in agreement with the ground truth. The SAEM-estimated locations of the two active sources are near the true locations while the extra (quiet) sources are localized in the opposite hemisphere. The chord plot of the gPDC matrix reveals bi-directional connectivity between the sources in the right lateral occipital (Source 3) and right fusiform (Source 5) parcel of the Desikan–Killiany (DK) atlas, with a stronger connection from Source 3 to Source 5 (most prominent in the bands of 4 − 8 Hz and 8 − 13 Hz). However, there is also a spurious connection between the extra sources: Source 1 → Source 2. The estimates of the gPDC matrix obtained with the SAEM correspond well with the ground truth as shown in Figure 6(A).

**Figure 6.**
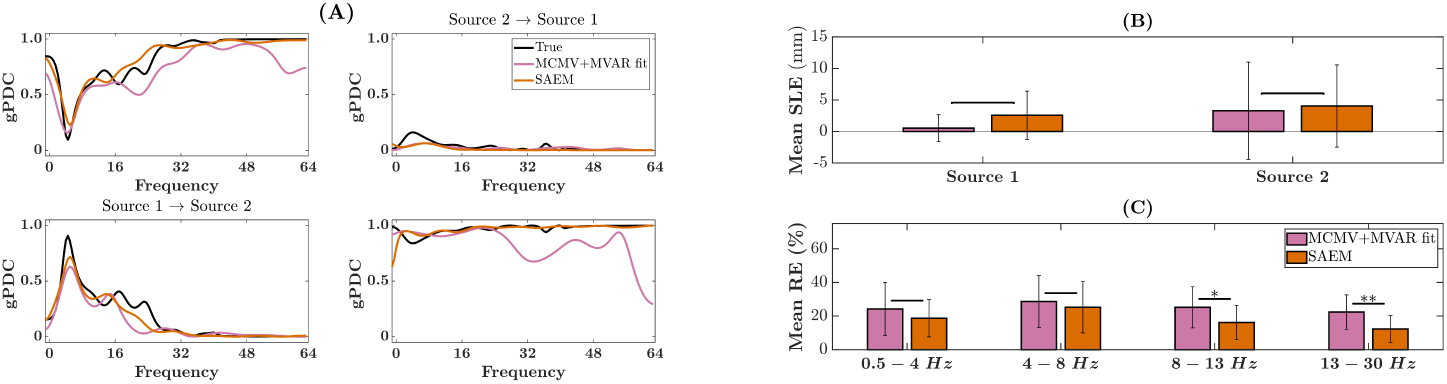
ECoG-based MEG simulations. (A) True, SAEM-estimated and MCMV+MVAR-estimated connectivity between Source 1 (within the right lateral occipital parcel of the DK atlas) and Source 2 (within the right fusiform gyrus parcel of the DK atlas) as a function of frequency. (B) Source localization error (SLE) of SAEM and MCMV methods for Sources 1 and 2 across the 16 subjects. (C) Mean reconstruction error (RE) of the gPDC matrix across the 16 subjects for both SAEM and MCMV+MVAR methods. ^*^*p* < 0.05, ^**^*p* < 0.01 and ^***^*p* < 0.001.

When applying the MCMV method followed by MVAR fitting (see Figure 5), the estimated amplitudes of two sources (labeled Source 1 and 2 in this case) correspond well to the ground truth but one of the extra sources, Source 3, also has a considerable amplitude. The other two extra sources (Sources 4 and 5) have amplitudes close to zero as they should. The estimated locations of Sources 1 and 2 are in a good agreement with the ground truth. The gPDC matrix obtained by MVAR fitting to source amplitudes shows bi-directional connectivity between Source 2 (in the right lateral occipital parcel of the DK atlas) and Source 1 (in the right fusiform parcel of the DK atlas), again with a stronger connection from the superior to inferior source. However, there are several spurious connections due to the extra sources that were estimated to be active (Source 4 → Source 2, Source 3 → Source 2, Source 5 → Source 1). Figure 6(A) also shows that MCMV followed by MVAR fitting underestimates the gPDC (in bands of 4 − 8 Hz and 8 − 13 Hz) in comparison to the gPDC estimate obtained with the SAEM method.

Figure 6(B) shows a quantitative comparison of SLE across the group of 16 subjects; the MCMV and SAEM methods perform similarly (no statistically significant difference between the mean SLEs). As seen in Figure 6 (C), both the two-step approach and the proposed SAEM method give similar RE of the gPDC matrix in the 4 − 8 Hz, where the connectivity between the true sources is the strongest. However, the SAEM method gives lower RE of the gPDC matrix compared to MCMV followed by MVAR fitting (*p* < 0.05) within 8 − 13 Hz and 13 − 30 Hz where the true connectivity between these sources is still significant, but not as dominant as in the 4 − 8 Hz band. Figures S8–S9 in the supplementary material show these results for individual subjects.

### 3.3 Real MEG data

We applied our SAEM algorithm to real MEG data recorded while the participants were presented with images of human faces [40]. Similarly as in the ECoG-based simulations, we assumed the number of sources to be 5.

#### 3.3.1 Single-subject analysis

Figure 7 shows the estimates of the source dipole locations (averaged over time), amplitudes and gPDC for a representative subject (SUB-10). The most active sources were localized to the left pericalcarine cortex (Source 5), peaking at around 120 ms after the stimulus image onset, to the left and right inferior temporal cortex (Source 3 and Source 1), peaking at 100–120 ms and 150–170 ms respectively to parts of right cingulate (Source 3). The source localized to orbitofrontal cortex (Source 4) had considerably lower amplitude. The directed functional connectivity as given by gPDC showed a prominent connection in the direction Source 5 (pericalcarine cortex) → Source 2 (inferior temporal cortex). Although gPDC was significant in all the four frequency bands, higher gPDC values were obtained within 0.5 − 4 Hz and 4 − 8 Hz. Significant but weaker interaction was also seen in the opposite direction. Bidirectional connections were also observed between left and right infereior temoporal cortices, with prominent connection Source 1 → Source 2.

**Figure 7.**
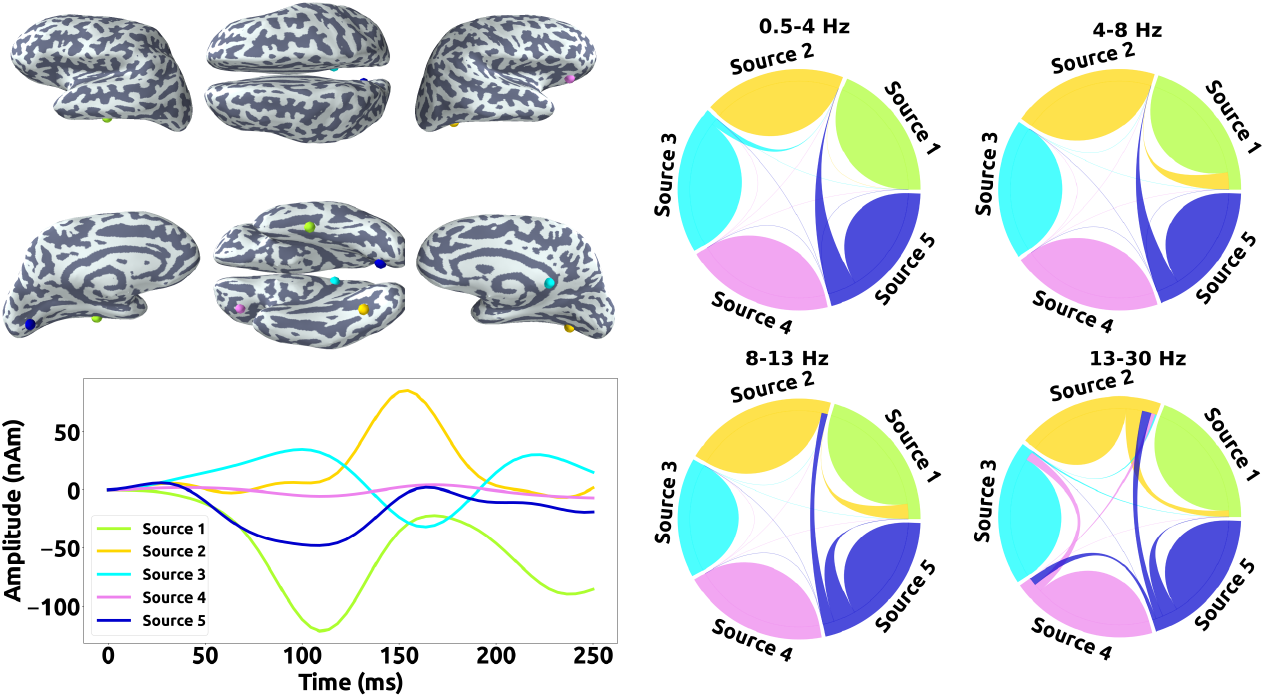
Real MEG data. SAEM-estimated source locations, amplitudes and functional connections in one subject.

#### 3.3.2 Group analysis

After estimating the source and connectivity parameters individually for all the subjects, we aggregated group-level results using clustering. To this end, we morphed the source locations of each subject to an average brain (‘fsaverage’ of FreeSurfer). We then clustered the sources using hierarchical clustering algorithm, with distance threshold. The cluster centroids, given as the spatial mean of source locations within each cluster, are shown in Figure 8. The centroids are located in the following regions of the Desikan–Killiany atlas: left lingual gyrus (LG -lh), left parahippocampal gyrus (PHG-lh), right parahippocampal gyrus (PHG-lh), right fusiform gyrus (FG-rh), left transverse temporal cortex (TTC-lh), right caudal anterior cingulate cortex (CAAC-rh), left isthumus cingulate cortex (ICC-lh), right lateral orbitofrontal cortex (LOFC-lh) and right postcentral gyrus (PG-rh).

**Figure 8.**
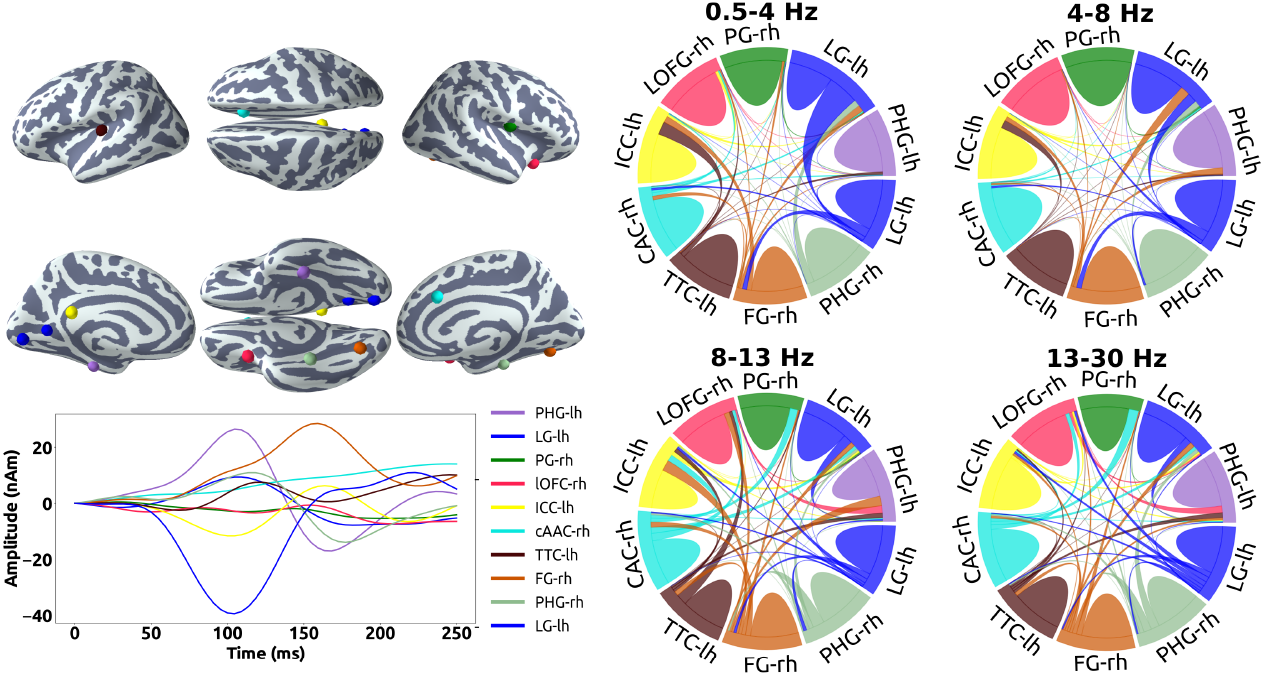
Real MEG data. Group-level (*N* = 16) estimates of source locations, amplitudes and their directed functional connections. Indicated locations are the source-cluster centroids, time courses are averages of those of individual sources in each cluster, and connectivity plots are derived from the gPDC matrices averaged across subjects.

Figure 8 also shows the combined amplitude of the sources (using the “flipped-mean” method) within each cluster. Sources in the occipital and inferior temporal cortex again exhibit distinct peaks at 120 and 170 ms. Finally, the directed functional connectivity as given by gPDC at the group level was obtained by averaging the entries of the subject-specific MVAR matrices pertaining to sources within each cluster. The strongest connectivity was again observed between occipital and inferior temporal areas (Sources in LG-lh ↔ FG-rh). Although the connection was bidirectional, stronger flow was observed from occipital (LG-lh) to ventral parts of the temporal region (FG-rh).

## 4 Discussion

In the present study, we propose a method to jointly estimate the locations and amplitudes of neural sources and their functional connections from MEG data. We use a conditionally linear state-space formulation, where the time-delayed correlation between the sources is given by a multivariate autoregressive (MVAR) model, whose spectral representation can reveal directed interactions between the sources at specific frequencies. Given the number of sources, our algorithm uses a conditional particle filter with ancestor sampling to estimate the source locations and a Kalman smoother to estimate the source amplitudes. The MVAR matrix and the covariance matrices of process and measurement noise are then obtained as the maximum-likelihood estimates within the stochastic approximation expectation–maximization (SAEM) framework. Using MVAR simulations of a simple source structure as well as more realistic ECoG-based simulations of MEG measurements, we show that the proposed joint estimation method outperforms the traditional two-step approach used for connectivity estimation. We also applied our method to real MEG data containing responses to visually-presented faces.

Fukushima and colleagues, driven by a similar motivation as ours, have developed a joint estimation method to estimate functional connectivity using a dynamic variational Bayesian approach [12]. However, their method still requires an anatomical prior in the form structural connectivity to constrain the MVAR matrix; such prior information may not always be available or even reliable. Dynamical causal modelling [10] is another hypothesis-driven approach that requires prior specification of source locations and their interaction structure, which also may not always be known. In contrast to these two approaches, our method only assumes that the MEG data can be explained by a small number of sources (< 5) – modeled as current dipoles – and jointly estimates the location, amplitude and connectivity between these sources without requiring any anatomical constraint or prior specification of source locations or ROIs.

### 4.1 Source estimation

The accuracy of estimating the locations of the sources underlying MEG responses naturally depends on the signal-to-noise ratio (SNR) of the data. Unaveraged MEG responses are typically considered to have an SNR of about 3–5; a study on MEG responses to faces estimated an empirical SNR of 4.0–4.3 [19]. When SNR_brain_ *≥* 3.0 in the MVAR simulations, our SAEM algorithm recovered more than 80% of the sources with less than 20-mm error (see Table 1). Thus, our algorithm should show at least decent source-localization accuracy already for unaveraged responses and in task-based MEG the accuracy shall improve with trial averaging, which the algorithm is designed to benefit from.

With the ECoG-based simulations presented in this paper, the SAEM algorithm estimated sources using an average of 40 trials. Despite the introduced modelling error and low SNR_brain_, the SAEM method gave accurate localization results for all the 16 subjects (2.5±3.8mm for the source in the right lateral occipital region and 4.0±6.5mm for the source in the right fusiform gyrus parcel of the DK atlas). Regarding source localization, we did not find a significant difference between the MCMV and SAEM algorithms. It has been previously shown that MCMV-based algorithms give reliable source estimates, even with arbitrary noise covariance matrices [29].

There has recently been interest in exploiting MCMV beamformers for estimating connectivity in MEG data [30]. Similarly to our SAEM, the MCMV approach assumes that a few point-like sources (*N*_*s*_ < 8) are responsible for generating the MEG data [30] and the number of these sources must be specified a priori. If the given number is higher than the true number of sources, the extra sources should ideally be estimated to have close-to-zero amplitude. We found that SAEM indeed shows baseline-noise-level amplitudes for these extra sources while the MCMV algorithm yielded considerably higher amplitudes, leading to spurious connections between the extra and true sources as seen in Figure 5 and in Figures S8–S9 in the Supplementary material.

### 4.2 Connectivity estimates

We employed simple MVAR-based simulations to compare the four algorithms in their ability to estimate functional connectivity. In these comparisons, the three two-step methods were given the advantage of knowing the source locations while SAEM estimated them from the data.

In the case of non-interacting sources (Type-0 scenario), the LCMV method followed by MVAR fitting outperformed the other methods as that scenario fulfilled the assumption of uncorrelated sources that LCMV makes. However, the SAEM method gave significantly less false positives than the LCMV method, despite not making any implicit assumptions about the nature of interaction between the sources (Type-I scenario).

However, when the sources were interacting, the SAEM method clearly outperformed the three two-step approaches used in this work. The performance difference is highlighted in the false positive rate for interacting sources: while the two-step approaches gave a high number of false connections, the SAEM method yielded significantly less, even at low SNR_brain_. Among the two-step approaches tested, MCMV beamforming followed by MVAR fitting outperformed the LCMV and eLORETA methods, which was expected since the MCMV method has been shown to be insensitive to source correlations [29].

SAEM utilizes maximum likelihood (ML) estimation of connectivity parameters, and it is known that ML gives asymptotically unbiased estimates with increasing sample size. In addition, since we also estimate process noise covariance explicitly, our approach leads to more accurate reconstruction of the MVAR matrix and consequently the gPDC matrix. In contrast, as shown in Ref. [4], the two-step approaches try to fit the MVAR model to the estimated source signals without taking into account the noise, which leads to biased MVAR estimates, particularly at low SNRs. The MCMV approach, despite being less affected by source correlations, yields biased source amplitudes for this reason, which eventually affects connectivity estimation. It is common in two-step MVAR modelling of connectivity to pre-select the locations of the sources or ROIs. Despite making no such assumption, our method gives connectivity estimates which are similar and in many cases superior in accuracy to those obtained with two-step approaches for correlated sources.

### 4.3 Application to real data

We applied SAEM to real MEG data recorded while the subjects were viewing still images of human faces. SAEM estimated sources in the occipital, temporal and cingulate cortices, which are the brain regions typically associated with adult face processing (see e.g. Refs [15, 17]). In addition, these locations are in agreement with previous MEG/EEG studies that have utilized either dipole localization [21, 34, 41] or distributed source imaging methods [28]. They were also largely consistent across the 16 subjects as was also revealed by the group analysis. The time courses of the estimated sources peaked at around 120 ms and 170 ms, again in agreement with previous studies [21, 41].

Although the location of sources involved in face processing and the latencies of source waveforms have been extensively studied, it is still not clear how these regions interact. Studies utilizing fMRI have reported correlations between occipital face area (OFA), fusiform face area (FFA) and superior temporal sulcus (STS) [5], whereas directed interaction from OFA to FFA and from FFA to STS has been reported [8]. The above-mentioned study by Fukushima and colleagues also reported significant interaction from OFA to FFA [12]. A recent study utilizing DCM on MEG data also found reciprocal connections between OFA and FFA [18]. Although a rigorous investigation of the existence of these connections and their dynamics is beyond the scope of this paper, we find that our results are largely in agreement with the results reported in literature regarding functional connectivity during face processing.

Our ECoG-based simulations were derived from real measurements that contained responses to viewing faces; the strongest responses were seen in the ECoG electrodes in superior and inferior regions in the right occipital lobe. In line with the studies discussed above, an MVAR model fitted to these two signals revealed bidirectional interaction, with the dominant direction being occipital → inferior temporal, which is in agreement with our SAEM results on real MEG data from a similar experiment: we observed bidirectional interactions between sources in the occipital and inferior temporal regions, with a stronger connection from occipital to inferior temporal.

### 4.4 Limitations

Since source localization in our algorithm is based on the likelihood function, in situations where there are two or more sources with very different strengths, our SAEM algorithm is likely to mislocalize or discard the weaker source(s). This problem is common to other source localization methods as well and is particularly pronounced in low SNR conditions. As seen from our results from MVAR simulations, the localization accuracy of the SAEM method improves with with the increasing SNR.

One of the limitations of our approach is that it requires specifying the number of sources, which may not always be known. If the number of sources set is higher than the actual number of sources, the algorithm localizes the extra sources to fit the noise, and the amplitudes of these extra sources will be close to zero, and thus they can be discarded. In case the given number of sources is less than the actual, the algorithm finds the strongest sources. In any case, we do not recommend setting the number of sources very high to keep the computational cost feasible as the Kalman filter scales cubically with the state dimension, which is *N*_*s*_*P*, where *N*_*s*_ is the number of sources and *P* is the model order. In order to run the algorithm in a reasonable time, we utilized GPUs, which reduced the computational time by a factor of 10–15 when compared to the CPU-only implementation. For *N*_*s*_ = 5, *N*_*p*_ = 500 *J* = 100 and *P* = 10, the algorithm takes approximately 60 s per SAEM iteration with NVIDIA® V100 Tensor Core GPU and requires 300–500 iterations to converge.

Another shortcoming is that we assume connectivity to be static, which is not necessarily true. We have previously proposed a method to estimate dynamic connectivity using a joint Kalman filter from MEG data [38]. However, this approach requires the source locations to be specified. Future work will be in the direction of estimating whole-brain dynamic functional connectivity, possibly applying methods based on variational Bayes or ensemble Kalman filters. Finally, although we use only MEG data in this work, it should be straightforward to apply the proposed algorithm to EEG data.

## 5 Conclusion

In this study, we proposed a novel framework based on the SAEM algorithm for joint estimation of source and connectivity parameters of neural sources from MEG data. Using both MVAR- and ECoG-based simulations, we demonstrated that the joint estimation framework provides more accurate reconstruction of functional connectivity coefficients compared to the state-of-the-art two-step approaches. Using visual-task-related MEG data from 16 subjects, we showed that our method gives physiologically plausible functional connectivity estimates both at individual and group level.

## 6 Data availability

All data used in this study are from open-access repositories and the corresponding links are provided in the text. The implementation of the SAEM algorithm, code for generating the simulated data and a script for pre-processing the MEG data are available at https://github.com/narayanps/SAEM_codes/tree/master.

## 7 Acknowledgements

This work was supported by Academy of Finland, grant BRAINTRACK (#289108) to L.P.

## Notes

https://legacy.openfmri.org/

https://github.com/narayanps/SAEM_codes/tree/master

### Competing Interest Statement

The authors have declared no competing interest.

### Summary of Updates

Added github link that includes all the code.

