## Supplementary text for "Joint estimation of neural sources and their functional connections from MEG data"

### 1 Methods

#### 1.1 State space model

We can re-write a $P$-th order MVAR model as a first-order model as follows:

$$\underbrace{\begin{bmatrix} q_t^{j,1} \\ \vdots \\ q_t^{j,n} \\ \vdots \\ q_{t-p+1}^{j,1} \\ \vdots \\ q_{t-p+1}^{j,n} \end{bmatrix}}_{\tilde{\mathbf{q}}_t^j} = \underbrace{\begin{bmatrix} a_{11,1} & \cdots & a_{1n,1} & \cdots & a_{11,p} & \cdots & a_{1n,p} \\ \vdots & \vdots & \vdots & \vdots & \vdots & \vdots & \\ a_{n1,1} & \cdots & a_{nn,1} & \cdots & a_{n1,p} & \cdots & a_{nn,p} \\ 1 & 0 & 0 & 0 & 0 & \cdots & 0 \\ 0 & \ddots & 0 & 0 & 0 & \cdots & 0 \\ 0 & 0 & \ddots & 0 & 0 & \cdots & 0 \\ 0 & 0 & 0 & 1 & 0 & \cdots & 0 \end{bmatrix}}_{\tilde{\mathbf{A}}} \underbrace{\begin{bmatrix} q_{t-1}^{j,1} \\ \vdots \\ q_{t-1}^{j,n} \\ \vdots \\ q_{t-p}^{j,1} \\ \vdots \\ q_{t-p}^{j,n} \end{bmatrix}}_{\tilde{\mathbf{q}}_{t-1}^j} + \underbrace{\begin{bmatrix} v_t^{j,1} \\ \vdots \\ v_t^{j,n} \\ \vdots \\ 0 \\ \vdots \\ 0 \end{bmatrix}}_{\tilde{\mathbf{v}}_t^j}. \quad (1)$$

The $np \times np$ covariance matrix, $\tilde{\mathbf{V}}$, of process noise $\tilde{\mathbf{v}}_t^j$ is given as

$$\begin{bmatrix} var(v^{j,1}, v^{j,1}) & \cdots & cov(v^{j,1}, v^{j,N_s}) & 0 & \cdots & 0 \\ \vdots & \vdots & \vdots & \vdots & \vdots & \\ cov(v^{j,N_s}, v^{j,1}) & \cdots & var(v^{j,N_s}, v^{j,N_s}) & 0 & \cdots & 0 \\ 0 & \cdots & 0 & 0 & \cdots & 0 \\ \vdots & \vdots & \vdots & \vdots & \vdots & \\ 0 & \cdots & 0 & 0 & \cdots & 0 \end{bmatrix}. \quad (2)$$

#### 1.2 Conditional particle filter with ancestor sampling

Before introducing conditional particle filter with ancestor sampling (CPF-AS), we will briefly review sequential Monte-Carlo (SMC) technique and then build on the concept of CPF-AS. In order to give a brief introduction to the standard SMC technique, also known as particle filter, we will consider a general nonlinear SSM of the form

$$\begin{aligned} \mathbf{x}_t &\sim p_\theta(\mathbf{x}_t|\mathbf{x}_{t-1}) \\ \mathbf{y}_t &\sim g_\theta(\mathbf{y}_t|\mathbf{x}_t). \end{aligned} \quad (3)$$

Using a particle filter, a sequence of target distributions $p_{\boldsymbol{\theta}}(\mathbf{x}_{1:t-1}|\mathbf{y}_{1:t-1})$ can be approximated as [4]

$$\widehat{p}_{\boldsymbol{\theta}}(\mathbf{dx}_{1:t-1}|\mathbf{y}_{1:t}) = \sum_{i=1}^{N_p} w_{t-1}^i \delta_{\mathbf{x}_{1:t-1}^i}(\mathbf{dx}_{1:t-1}) \tag{4}$$

where $\{\mathbf{x}_{1:t-1}, w_{t-1}^i\}_{i=1}^{N_p}$ is a weighted particle system, with $\sum_i w_{t-1}^i = 1$. Using sequential importance resampling [6], the particles at time $t-1$ are propagated to time $t$. This entails sampling the ancestor index with

$$P(a_t^i = m) \propto w_{t-1}^m \tag{5}$$

for $i = \{1 \cdots N_p\}$. A new set of particles at time $t$ are then drawn according to

$$\mathbf{x}_t^i \sim q_{\boldsymbol{\theta}}(\mathbf{x}_t|\mathbf{x}_{t-1}^{a_t^i}, \mathbf{y}_t) \tag{6}$$

where $q_{\boldsymbol{\theta}}(\cdot)$ is some suitable proposal kernel. The particle trajectories can now be extended as $\mathbf{x}_{1:t} := \{\mathbf{x}_{1:t-1}^{a_t^i}, \mathbf{x}_t\}$. The normalized weights $w_t^i$ are given as

$$w_t^i \propto \frac{p_{\boldsymbol{\theta}}(\mathbf{y}_t, \mathbf{x}_t|\mathbf{y}_{1:t-1}, \mathbf{x}_{1:t-1})}{q_{\boldsymbol{\theta}}(\mathbf{x}_t|\mathbf{y}_{1:t}, \mathbf{x}_{1:t-1})}. \tag{7}$$

In the CPF-AS method, we set the $N_p$-th particle trajectory deterministically, i.e., $\mathbf{x}_{1:T}^{N_p} = \mathbf{x}'_{1:T}$ and the sampling given in Equation 6 is done only for particles $i = 1 \dots N_p - 1$. The ancestor index $a_t^{N_p}$ for the $N_p$-th particle is sampled according to

$$P(a_t^{N_p} = m) \propto w_{t-1}^m p_{\boldsymbol{\theta}}(\mathbf{x}'_t|\mathbf{x}_{t-1}^m). \tag{8}$$

The CPF-AS is in fact a modification of particle Gibbs (PG) algorithm [1]. Thus, the CPF-AS is similar to a particle filter except that the trajectory of one particle, known as reference trajectory, is fixed *a priori* [3, 4].

Note that one could also use a particle smoother in the EM algorithm to estimate $p_\theta(\mathbf{x}_{1:T}|\mathbf{y}_{1:T})$. However, this entails a computational complexity of $\mathcal{O}(N_p^2 KT)$, where $N_p$ is the number of particles and $K$ is the number of EM iterations and $T$ is the number of time samples. Using PMCMC with particle filter as the kernel reduces the computational complexity to $\mathcal{O}(N_p KT)$.

In our model, $\mathbf{x}_{1:T} = \{\mathbf{r}_{1:T}, \tilde{\mathbf{q}}_{1:T}\}$. Due to Rao–Blackwellization, we need to use the CPF-AS only to approximate $p_\theta(\mathbf{r}_{1:T}|\bar{\mathbf{y}}_{1:T})$. We sample the particles, i.e., the source locations at time $t$, according to the model $\mathbf{r}_t^i \sim \mathcal{N}(\mathbf{m}_{t-1}^i, h^2 \boldsymbol{\Xi}_{t-1})$ with

$$\mathbf{m}_{t-1}^i = a\mathbf{r}_{t-1}^{a_t^i} + (1-a)\bar{\mathbf{r}}_{t-1} \tag{9}$$

where $\bar{\mathbf{r}}_{t-1}$ and $\boldsymbol{\Xi}_{t-1}$ are the mean and variance of the Monte-Carlo approximation to $p_\theta(\mathbf{r}_{t-1}|\bar{\mathbf{y}}_{1:t-1})$. We first set the discount factor $\delta \in (0, 1]$ and compute the tuning parameter $a = (3\delta - 1)/2\delta$ and the smoothing parameter $h^2 = 1 - a^2$. In this work, we set $\delta = 0.95$.

Due to the non-Markovian nature of $\mathbf{r}_{1:T}|\bar{\mathbf{y}}_{1:T}$, with $\tilde{\mathbf{q}}_{1:T}$ marginalized out [3, 7], the ancestor index does not have the simple expression as given in Equation 8. Instead, for our model, the ancestor index of particle $N_p$ is sampled according to $P(a_t^{N_p} = i) \propto w_{t-1}^i p(\bar{\mathbf{y}}_{t:T}, \mathbf{r}'_{t:T}|\mathbf{r}_{1:t-1}^i, \bar{\mathbf{y}}_{1:t-1})$, where $\bar{\mathbf{y}}_t = \sum_{j=1}^J \mathbf{y}_t^j$. Using the notation in Ref. [3], we define $||\boldsymbol{\mu}_{t|1:t}||_{\boldsymbol{\Omega}}^2 \triangleq \boldsymbol{\mu}_{t|1:t}^T \boldsymbol{\Omega} \boldsymbol{\mu}_{t|1:t}$, $\mathbf{P}_{t|1:t} \triangleq \boldsymbol{\Gamma}_{t|1:t} \boldsymbol{\Gamma}_{t|1:t}^T$ and $\mathbf{V} \triangleq \mathbf{F}\mathbf{F}^T$. Furthermore, let us define $\tilde{\mathbf{F}} \in \mathbb{R}^{N_s P \times N_s P}$ as

$$\tilde{\mathbf{F}} = \begin{bmatrix} \mathbf{F} & \mathbf{0} \\ \mathbf{0} & \mathbf{0} \end{bmatrix} \tag{10}$$

and $\tilde{\mathbf{G}} \in \mathbb{R}^{M \times N_s P}$ as

$$\tilde{\mathbf{G}}_t = \begin{bmatrix} \mathbf{G}(\mathbf{r}_t^{N_p}) & \mathbf{0} \end{bmatrix}. \tag{11}$$

Finally, we have

$$p(\bar{\mathbf{y}}_{t:T}, \mathbf{r}'_{t:T} | \mathbf{r}^i_{1:t-1}, \mathbf{y}_{1:t-1}) \propto \boldsymbol{\Delta}^i_{t-1} |\boldsymbol{\Gamma}^i_{t-1}|^{-1/2} \exp(-\frac{1}{2}\boldsymbol{\eta}^i_{t-1}) \tag{12}$$

with

$$\begin{aligned} \boldsymbol{\Delta}^i_t &= p(\mathbf{r}'_t | \mathbf{r}^i_{t-1}) \\ \boldsymbol{\eta}^i_t &= ||\bar{\boldsymbol{\mu}}^i_{t|1:t}||^2_{\boldsymbol{\Omega}_t} - 2\bar{\boldsymbol{\lambda}}_t \bar{\boldsymbol{\mu}}^i_{t|1:t} - ||\bar{\boldsymbol{\Gamma}}^i_{t|1:t}(\bar{\boldsymbol{\lambda}}_t - \boldsymbol{\Omega}_t \bar{\boldsymbol{\mu}}^i_{t|1:t}||^2_{\boldsymbol{\Lambda}_t^{-1}} \\ \boldsymbol{\Lambda}^i_t &= \bar{\boldsymbol{\Gamma}}^{i,T}_{t|1:t} \boldsymbol{\Omega}_t \bar{\boldsymbol{\Gamma}}^i_{t|1:t} + \mathbf{I} \end{aligned} \tag{13}$$

where $\bar{\boldsymbol{\Gamma}}^i_{t|1:t} = \frac{1}{J}\sum_{j=1}^J \boldsymbol{\Gamma}^{i,j}_{t|1:t}$ and $\bar{\boldsymbol{\mu}}^i_{t|1:t}$, $\bar{\boldsymbol{\lambda}}_t$ is defined analogously. For $t = T-1 \ldots 1$, we define the following backward statistics:

$$\begin{aligned} \boldsymbol{\Omega}_t &= \tilde{\mathbf{A}}^T (\mathbf{I} - \widehat{\boldsymbol{\Omega}}_{t+1} \tilde{\mathbf{F}} \mathbf{M}^{-1}_{t+1} \tilde{\mathbf{F}}^T) \widehat{\boldsymbol{\Omega}}_{t+1} \tilde{\mathbf{A}}^T \\ \boldsymbol{\lambda}^j_t &= \tilde{\mathbf{A}}^T (\mathbf{I} - \widehat{\boldsymbol{\Omega}}_{t+1} \tilde{\mathbf{F}} \mathbf{M}^{-1}_{t+1} \tilde{\mathbf{F}}^T) \widehat{\boldsymbol{\lambda}}^j_{t+1} \\ \widehat{\boldsymbol{\Omega}}_t &= \boldsymbol{\Omega}_t + \tilde{\mathbf{G}}^T_t \mathbf{E}^{-1} \tilde{\mathbf{G}}_t \\ \widehat{\boldsymbol{\lambda}}^j_t &= \boldsymbol{\lambda}^j_t + \tilde{\mathbf{G}}_t \mathbf{E}^{-1} \mathbf{y}^j_t \\ \mathbf{M}_{t+1} &= \tilde{\mathbf{F}}^T \widehat{\boldsymbol{\Omega}}_{t+1} \tilde{\mathbf{F}} + \mathbf{I} \end{aligned} \tag{14}$$

with $\widehat{\boldsymbol{\Omega}}_T$ and $\widehat{\boldsymbol{\lambda}}_T$ given as

$$\begin{aligned} \widehat{\boldsymbol{\Omega}}_T &= \tilde{\mathbf{G}}^T_T \mathbf{E}^{-1} \tilde{\mathbf{G}}_T \\ \widehat{\boldsymbol{\lambda}}^j_T &= \tilde{\mathbf{G}}_T \mathbf{E}^{-1} \mathbf{y}^j_T. \end{aligned} \tag{15}$$

#### 1.3 Estimating source amplitudes – Kalman filter

For the $i$-th particle and $j$-th trial, the predictive mean and covariance of the Kalman filter are given as

$$\begin{aligned} \boldsymbol{\mu}^{i,j}_{t|1:t-1} &= \tilde{\mathbf{A}} \boldsymbol{\mu}^{i,j}_{t-1|1:t-1} \\ \mathbf{P}^{i,j}_{t|1:t-1} &= \tilde{\mathbf{A}} \mathbf{P}^{i,j}_{t-1|1:t-1} \tilde{\mathbf{A}}^T + \tilde{\mathbf{V}}. \end{aligned} \tag{16}$$

For the $i$-th particle and $j$-th trial, the updated mean and covariance of the Kalman filter, $\boldsymbol{\mu}^{i,j}_{t|1:t}$ and $\mathbf{P}^{i,j}_{t|1:t}$, are given as

$$\begin{aligned} \boldsymbol{\mu}^{i,j}_{t|1:t} &= \boldsymbol{\mu}^{i,j}_{t|1:t-1} + \mathbf{K}^{i,j}_t (\mathbf{y}^j_t - \mathbf{G}(\mathbf{r}^i_t) \boldsymbol{\mu}^{i,j}_{t|1:t-1}) \\ \mathbf{P}^{i,j}_{t|1:t} &= \mathbf{P}^{i,j}_{t|1:t-1} - \mathbf{K}^{i,j}_t \mathbf{S}^{i,j}_t (\mathbf{K}^{i,j}_t)^T \\ \mathbf{S}^{i,j}_t &= \mathbf{G}(\mathbf{r}^i_t) \mathbf{P}^{i,j}_{t-1|t-1} \mathbf{G}(\mathbf{r}^i_t)^T + \mathbf{E} \\ \mathbf{K}^{i,j}_t &= \mathbf{P}^{i,j}_{t|t-1} \mathbf{G}(\mathbf{r}^i_t)^T (\mathbf{S}^{i,j}_t)^{-1}. \end{aligned} \tag{17}$$

For the $i$-th particle and $j$-th trial, the mean and covariance of the Kalman smoother, $\boldsymbol{\mu}^{i,j}_{t|1:T}$ and $\mathbf{P}^{i,j}_{t|1:T}$ for $t = T-1 \ldots 1$, are given as

$$\begin{aligned} \boldsymbol{\mu}^{i,j}_{t|1:T} &= \boldsymbol{\mu}^{i,j}_{t|1:t} + \mathbf{J}^{i,j}_t (\boldsymbol{\mu}^{i,j}_{t|1:T+1} - \boldsymbol{\mu}^{i,j}_{t|1:t-1}) \\ \mathbf{J}^{i,j}_t &= \mathbf{P}^{i,j}_{t|1:t} \tilde{\mathbf{A}}^T (\mathbf{P}^{i,j}_{t|1:t-1})^{-1} \\ \mathbf{P}^{i,j}_{t|1:T} &= \mathbf{P}^{i,j}_{t|1:t} + \mathbf{J}^{i,j}_t (\mathbf{P}^{i,j}_{t|1:T+1} - \mathbf{P}^{i,j}_{t|1:t-1}) (\mathbf{J}^{i,j}_t)^T. \end{aligned} \tag{18}$$

Finally, the one-lag covariance smoother is given as

$$\mathbf{P}^{i,j}_{t,t-1|1:T} = \mathbf{P}^{i,j}_{t|1:t} (\mathbf{J}^{i,j}_{t-1})^T + \mathbf{J}^{i,j}_t (\mathbf{P}^{i,j}_{t+1,t|1:T} - \tilde{\mathbf{A}} \mathbf{P}^{i,j}_{t|1:t}) (\mathbf{J}^{i,j}_{t-1})^T. \tag{19}$$

#### 1.4 ML estimation of parameters

##### 1.4.1 Update for $\tilde{\mathbf{A}}$

At iteration $k$, to maximize $\hat{\mathcal{Q}}_k(\boldsymbol{\theta})$ w.r.t $\tilde{\mathbf{A}}$, we set $\frac{\partial \mathcal{Q}_k(\boldsymbol{\theta})}{\partial \tilde{\mathbf{A}}} = 0$. Separating only the terms involving $\tilde{\mathbf{A}}$ in $\hat{\mathcal{Q}}_k(\boldsymbol{\theta})$ and using Equations 103, 104 and 118 in Ref. [5] we obtain

$$\begin{aligned} \frac{\partial \mathcal{Q}_k(\boldsymbol{\theta})}{\partial \tilde{\mathbf{A}}} &= -\frac{1}{2}[-\frac{\partial}{\partial \tilde{\mathbf{A}}} tr(\tilde{\mathbf{V}}^{-1}\boldsymbol{\Psi}\tilde{\mathbf{A}}^T) - \frac{\partial}{\partial \tilde{\mathbf{A}}} tr(\tilde{\mathbf{V}}^{-1}\tilde{\mathbf{A}}\boldsymbol{\Psi}^T) + \frac{\partial}{\partial \tilde{\mathbf{A}}} tr(\tilde{\mathbf{V}}^{-1}\tilde{\mathbf{A}}\boldsymbol{\Sigma}\tilde{\mathbf{A}}^T)] \\ &= \tilde{\mathbf{V}}^{-1}\boldsymbol{\Psi} - \tilde{\mathbf{V}}^{-1}\tilde{\mathbf{A}}\boldsymbol{\Sigma}^{\boldsymbol{T}}. \end{aligned} \tag{20}$$

Since $\boldsymbol{\Sigma}$ is symmetric, setting the above derivative to zero we obtain $\tilde{\mathbf{A}}_{k+1} = \boldsymbol{\Psi}\boldsymbol{\Sigma}^{-1}$.

##### 1.4.2 Update for $\tilde{\mathbf{V}}$

Again, separating the terms involving only $\tilde{\mathbf{A}}$ in $\hat{\mathcal{Q}}_k(\boldsymbol{\theta})$ and using Equations 57 and 124 in Ref. [5], we get

$$\begin{aligned} \frac{\partial \mathcal{Q}_k(\boldsymbol{\theta})}{\partial \tilde{\mathbf{V}}} &= -\frac{JT}{2}\frac{\partial}{\partial \tilde{\mathbf{V}}} \log|2\pi\tilde{\mathbf{V}}| - \frac{1}{2}[\frac{\partial}{\partial \tilde{\mathbf{V}}} tr(\tilde{\mathbf{V}}^{-1}\boldsymbol{\Phi}) - \frac{\partial}{\partial \tilde{\mathbf{V}}} tr(\tilde{\mathbf{V}}^{-1}\boldsymbol{\Psi}\tilde{\mathbf{A}}^T) - \frac{\partial}{\partial \tilde{\mathbf{V}}} tr(\tilde{\mathbf{V}}^{-1}\tilde{\mathbf{A}}\boldsymbol{\Psi}^T) + \text{[illegible]} \\ &= -\frac{JT}{2}\tilde{\mathbf{V}}^{-1} + \frac{1}{2}\tilde{\mathbf{V}}^{-1}[\boldsymbol{\Phi} - \boldsymbol{\Psi}\tilde{\mathbf{A}}^T - \tilde{\mathbf{A}}\boldsymbol{\Psi}^T + \tilde{\mathbf{A}}\boldsymbol{\Sigma}\tilde{\mathbf{A}}^T]\tilde{\mathbf{V}}^{-1}. \end{aligned} \tag{21}$$

Setting the above derivative to zero, we obtain $\tilde{\mathbf{V}}_{k+1} = \frac{1}{JT}[\boldsymbol{\Phi} - \boldsymbol{\Psi}\tilde{\mathbf{A}}^T - \tilde{\mathbf{A}}\boldsymbol{\Psi}^T + \tilde{\mathbf{A}}\boldsymbol{\Sigma}\tilde{\mathbf{A}}^T]$.

##### 1.4.3 Update for $\sigma_m$

When we only have measurement noise, separating the terms involving only $\mathbf{E} = \sigma_m^2\mathbf{I}$ in $\hat{\mathcal{Q}}_k(\boldsymbol{\theta})$, we obtain

$$\begin{aligned} \frac{\partial \mathcal{Q}_k(\boldsymbol{\theta})}{\partial \sigma_m} &= -\frac{JT}{2}\frac{\partial}{\partial \sigma_m} \log|2\pi\sigma_m^2\mathbf{I}| - \frac{1}{2}[\frac{\partial}{\partial \sigma_m} tr(\sigma_m^{-2}[\mathbf{Z} - \boldsymbol{\Upsilon}\bar{\mathbf{G}}^T - \bar{\mathbf{G}}\boldsymbol{\Upsilon}^T + \bar{\mathbf{G}}\boldsymbol{\Phi}\bar{\mathbf{G}}^T]) \\ &= -JTM\frac{1}{\sigma_m} + \sigma_m^3 tr(\mathbf{Z} - \boldsymbol{\Upsilon}\bar{\mathbf{G}}^T - \bar{\mathbf{G}}\boldsymbol{\Upsilon}^T + \bar{\mathbf{G}}\boldsymbol{\Phi}\bar{\mathbf{G}}^T). \end{aligned} \tag{22}$$

Again, setting the above derivative to zero we get
$\sigma_{m,k+1}^2 = \frac{1}{JMT} tr(\mathbf{Z} - \boldsymbol{\Upsilon}\bar{\mathbf{G}}^T - \bar{\mathbf{G}}\boldsymbol{\Upsilon}^T + \bar{\mathbf{G}}\boldsymbol{\Phi}\bar{\mathbf{G}}^T)$, where $M$ is the number of MEG sensors.

##### 1.4.4 Update for $\sigma_b$ and $\sigma_m$

When biological noise is also included, the total noise covariance $\mathbf{R} = \sigma_m^2\mathbf{I} + \sigma_b^2\mathbf{G}\mathbf{G}^T$. Separating terms involving only $\sigma_b^2$ and $\sigma_m^2$, the update of $\sigma_b$ and $\sigma_m$ can be done using gradient descent with backtracking line search [2], where the goal is to minimize the objective function

$$\mathcal{L}_k(\boldsymbol{\theta}) = -\mathcal{Q}_k(\boldsymbol{\theta}) = \frac{JT}{2}\log|2\pi\mathbf{R}| + \frac{1}{2} tr\{\mathbf{R}^{-1}[\mathbf{Z} - \boldsymbol{\Upsilon}\bar{\mathbf{G}}^T - \bar{\mathbf{G}}\boldsymbol{\Upsilon}^T + \bar{\mathbf{G}}\boldsymbol{\Phi}\bar{\mathbf{G}}^T]\} \tag{23}$$

with the following gradients obtained using the chain rule: [5]

$$\begin{aligned} \frac{\partial \mathcal{L}_k(\boldsymbol{\theta})}{\partial \sigma_b} &= tr\{(\frac{\partial \mathcal{L}_k(\boldsymbol{\theta})}{\partial \mathbf{R}})^T \frac{\partial \mathbf{R}}{\partial \sigma_m}\} \\ &= tr\{(\mathbf{R}^{-1} - \mathbf{R}^{-1}[\mathbf{Z} - \boldsymbol{\Upsilon}\bar{\mathbf{G}}^T - \bar{\mathbf{G}}\boldsymbol{\Upsilon}^T + \bar{\mathbf{G}}\boldsymbol{\Phi}\bar{\mathbf{G}}^T]\mathbf{R}^{-1})^T(2\sigma_b\mathbf{G}\mathbf{G}^T)\} \end{aligned} \tag{24}$$

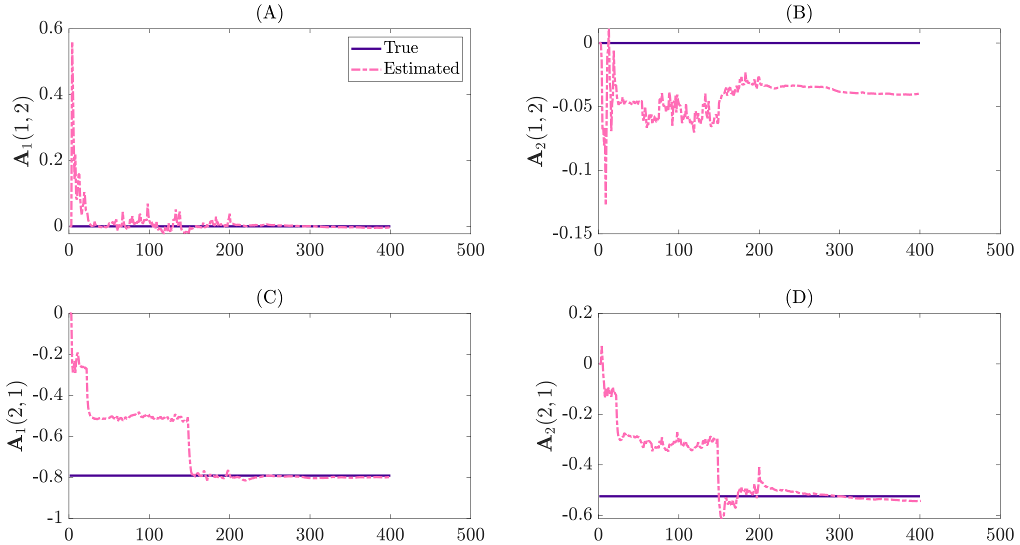


**Figure 1.** Estimates of the cross-coupling coefficients $\mathbf{A}_p(1,2)$ (A–B) and $\mathbf{A}_p(2,1)$ (C–D) for $p = 1$ to 2 obtained at every iteration using the SAEM algorithm (dashed lines) for an exemplary MVAR simulation. The true values are shown as solid lines.

and

$$\begin{aligned} \frac{\partial \mathcal{L}_k(\boldsymbol{\theta}))}{\partial \sigma_m} &= tr\{(\frac{\partial \mathcal{L}_k(\boldsymbol{\theta})}{\partial \mathbf{R}})^T \frac{\partial \mathbf{R}}{\partial \sigma_b}\} \\ &= tr\{(\mathbf{R}^{-1} - \mathbf{R}^{-1}[\mathbf{Z} - \boldsymbol{\Upsilon}\bar{\mathbf{G}}^T - \bar{\mathbf{G}}\boldsymbol{\Upsilon}^T + \bar{\mathbf{G}}\boldsymbol{\Phi}\bar{\mathbf{G}}^T]\mathbf{R}^{-1})^T(2\sigma_m\mathbf{I})\}. \end{aligned} \tag{25}$$

#### 2 Joint estimation algorithm and results

The joint estimation method is summarized in Algorithm 1 and the stochastic E-step used within the joint estimation scheme is presented in Algorithm 2.

**Algorithm 1** Joint estimation.

**procedure** SAEM
  Initialize $\boldsymbol{\theta}_0$ and $\mathbf{r}'_{1:T}[0]$ and $\hat{\mathcal{Q}}_0(\theta) = 0$
  **for** $k \geq 1$ to $K$ **do**
    Run RB-PMCMC to obtain $\{\mathbf{r}^i_{1:T}, w^i_T\}^N_{i=1}$, $\{\boldsymbol{\mu}^{i,j}_{1:T}\}^{i=N,j=J}_{i=1,j=1}$ and $\mathbf{r}_{1:T}[k]$.
    Compute $\hat{\mathcal{Q}}_k(\theta)$ in the stochastic E-step.
    Compute $\hat{\boldsymbol{\theta}}_k = \arg\max_\theta \hat{\mathcal{Q}}_k(\theta)$.
  **end for**
**end procedure**

**Algorithm 2** Stochastic E-step.

**procedure** RB-PMCMC
  **Input:** $\mathbf{y}_{1:T}$, $\bar{\mathbf{y}}_{1:T}$, $\mathbf{r}'_{1:T}[k]$
  **Output:** $\{\mathbf{r}^i_{1:T}, w^i_T\}_{i=1}^{N_p}$, $\{\boldsymbol{\mu}^{i,j}_{1:T}\}_{i=1,j=1}^{i=N_p,j=J}$ and $\mathbf{r}_{1:T}[k+1]$,
  $\mathbf{r}^i_t \sim p(\mathbf{r}_t)$ for $i = 1 \cdots N_p - 1$ and $t = 1$
  Compute $\boldsymbol{\Omega_t}$ and $\boldsymbol{\lambda_t}$ for $t = 1 \cdots T$ using Equation 14
  Set $\{\mathbf{r}^{N_p}_1 \cdots \mathbf{r}^{N_p}_T\} = \{\mathbf{r}'_1 \cdots \mathbf{r}'_T\}$
  Compute $w^i_t \propto p_{\boldsymbol{\theta}}(\bar{\mathbf{y}}_t|\mathbf{r}^i_t)$ for $i = 1 \cdots N_p$ and $t = 1$
  Compute $\boldsymbol{\mu}^{i,j}_{t|1:t}$ and $\boldsymbol{P}^{i,j}_{t|1:t}$ for $i = 1 \cdots N_p$, $j = 1 \cdots J$ and $t = 1$ Equation 17
  **for** $t = 2$ to $T$ **do**
    Draw $a^i_t$ with $\mathbb{P}(a^i_t = m) = w^m_{t-1}$ for $i = 1 \cdots N_p - 1$.
    $\mathbf{r}^i_t \sim p_{\boldsymbol{\theta}}(\mathbf{r}_t|\mathbf{r}^{a^i_t}_{t-1})$ for $i = 1 \cdots N_p - 1$.
    Compute $\boldsymbol{\Delta}^i_{t-1}$, $\boldsymbol{\eta}^i_{t-1}$, $\boldsymbol{\Lambda}^i_{t-1}$ for $i = 1 \cdots N_p$ using Equation 13.
    Draw $a^{N_p}_t$ according to $\mathbb{P}(a^{N_p}_t = i) \propto w^i_{t-1}\boldsymbol{\Delta}^i_{t-1}|\boldsymbol{\Lambda}^i_{t-1}|\exp(-\frac{1}{2}\boldsymbol{\eta}^i_{t-1})$.
    Set $\mathbf{r}^i_{1:t} = \{\mathbf{r}^{a^i_t}_{1:t-1}, \mathbf{r}^i_t\}$ for $i = 1 \cdots N_p$ .
    Set $\boldsymbol{\mu}^{i,j}_{1:t-1|1:t-1} = \boldsymbol{\mu}^{a^i_t,j}_{1:t-1|1:t-1}$ and $\boldsymbol{P}^{i,j}_{1:t-1|1:t-1} = \boldsymbol{P}^{a^i_t,j}_{1:t-1|1:t-1}$ for $i = 1 \cdots N_p$, $i = 1 \cdots N_p$ and $j = 1 \cdots J$.
    Set $\boldsymbol{\mu}^{i,j}_{1:t-1|1:t-2} = \boldsymbol{\mu}^{a^i_t,j}_{1:t-1|1:t-2}$ and $\boldsymbol{P}^{i,j}_{1:t-1|1:t-2} = \boldsymbol{P}^{a^i_t,j}_{1:t-1|1:t-2}$ for $i = 1 \cdots N_p$ and $j = 1 \cdots J$.
    Compute filtered mean $\boldsymbol{\mu}^{i,j}_{t|1:t}$ and covariance $\boldsymbol{P}^{i,j}_{t|1:t}$ for $i = 1 \cdots N_p$ and $j = 1 \cdots J$ using Equation 17.
    Compute predictive mean $\boldsymbol{\mu}^{i,j}_{t|1:t-1}$ and covariance $\boldsymbol{P}^{i,j}_{t|1:t-1}$ for $i = 1 \cdots N_p$ and $j = 1 \cdots J$ using Equation 16.
    $w^i_t \propto p_{\boldsymbol{\theta}}(\bar{\mathbf{y}}_t|\mathbf{r}^i_t, \bar{\mathbf{y}}_{1:t-1})$ for $i = 1 \cdots N_p - 1$.
    Normalize $w^i_t$ s.t. $\sum_i w^i_t = 1$.
  **end for**
  **for** $t = T$ to 1 **do**
    Compute smoothed mean $\boldsymbol{\mu}^{i,j}_{t|1:T}$ and covariance $\boldsymbol{P}^{i,j}_{t|1:T}$ for $i = 1 \cdots N_p$ and $j = 1 \cdots J$ using Equation 18.
  **end for**
  Set $\mathbf{r}[k+1] = \mathbf{r}^M_{1:T}$ with $\mathbb{P}(M = m) = w^m_T$.
**end procedure**

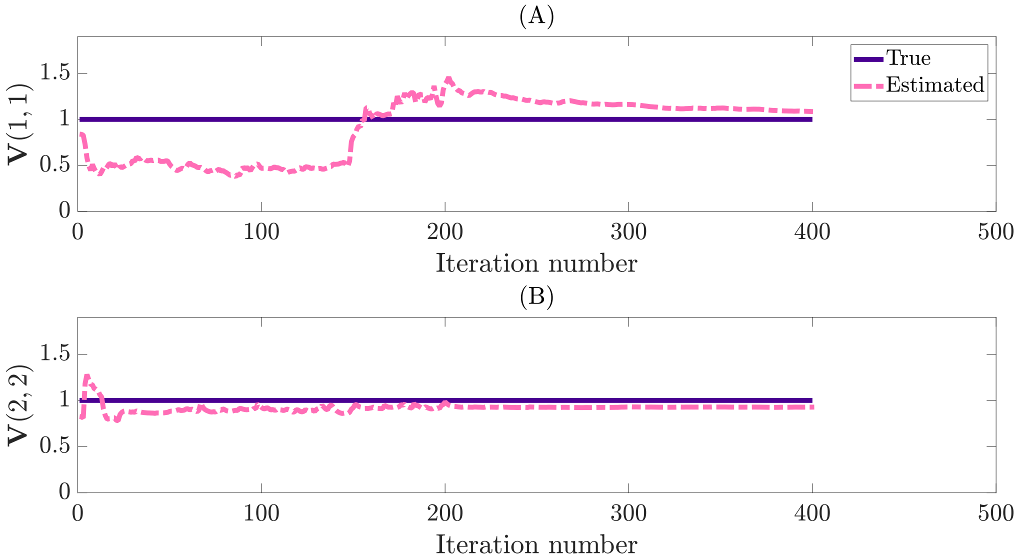


**Figure 2.** Estimate of the process noise covariance $\mathbf{V}(1,1)$ (A) and $\mathbf{V}(2,2)$ (B) obtained at every iteration using the SAEM algorithm (dashed lines) for an exemplary MVAR simulation. The true values are shown as solid lines.

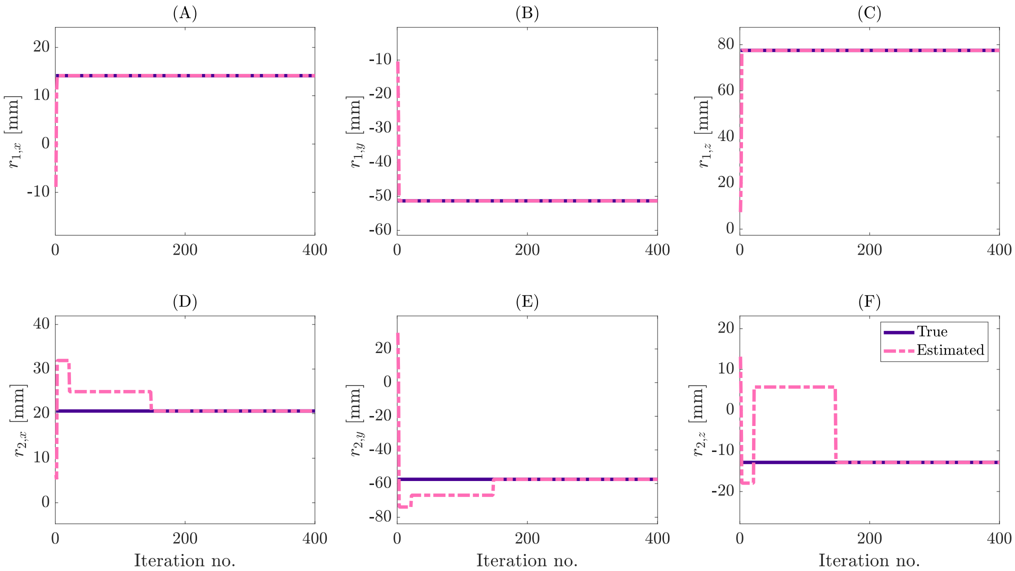


**Figure 3.** Estimate of the location ($r_x$, $r_y$, $r_z$) of Source 1 (A–C) and Source 2 (D–F) obtained at every iteration (at $t = T$) using the SAEM algorithm (dashed lines) for an exemplary MVAR simulation. The true values are shown as solid lines.

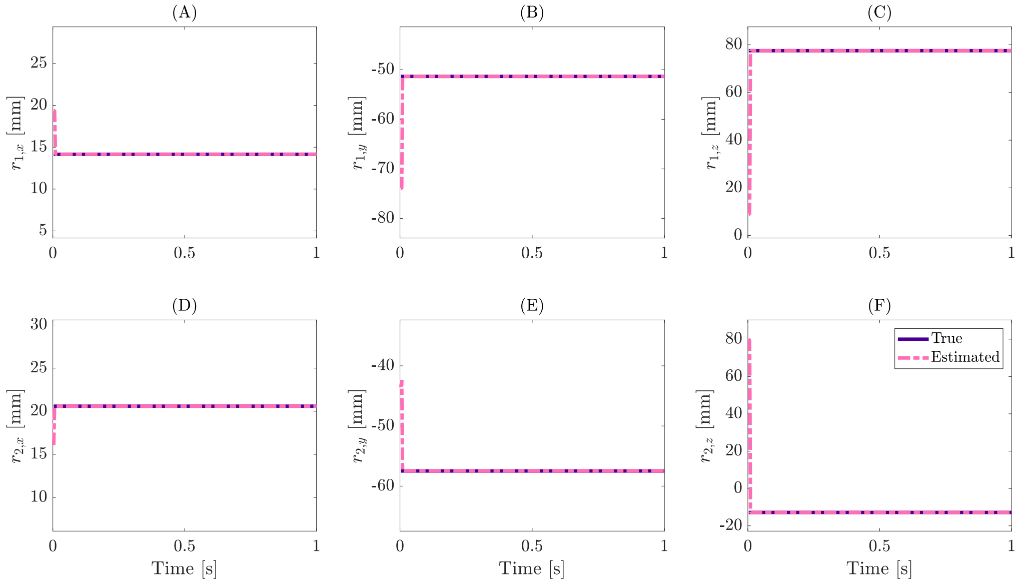


**Figure 4.** Converged estimate of the location ($r_x$, $r_y$, $r_z$) of Source 1 (A–C) and Source 2 (D–F) obtained using the SAEM algorithm (dashed lines) for an exemplary MVAR simulation. The true values are shown as solid lines. The estimate at every time point is given as the mean of the posterior distribution obtained from the PMCMC smoother.

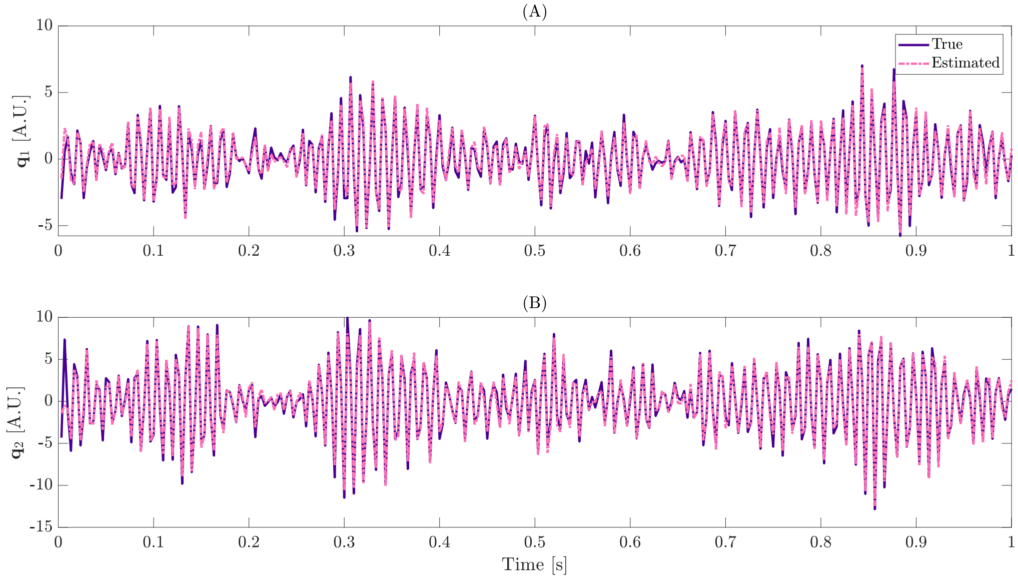


**Figure 5.** Converged estimate of the amplitude ($\mathbf{q}$) of Source 1 (A) and Source 2 (B) obtained using the SAEM algorithm (dashed lines) for an exemplary MVAR simulation. The true values are shown as solid lines.

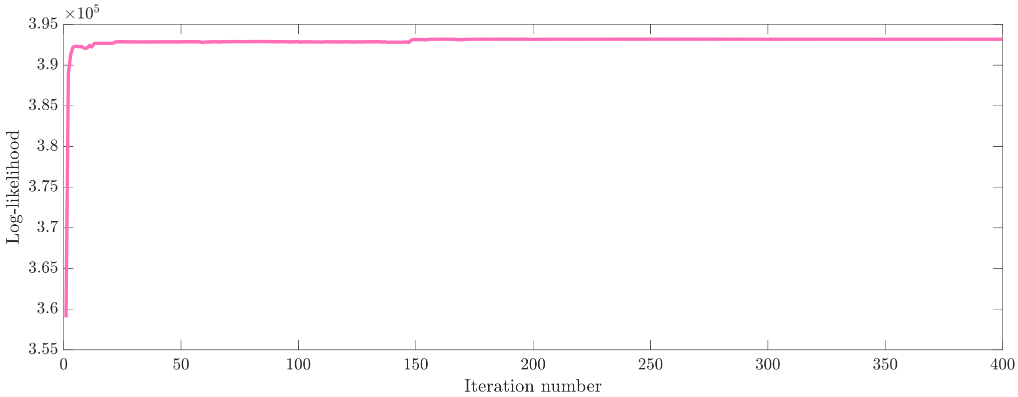


**Figure 6.** Complete data log-likelihood as a function of SAEM iterations for an exemplary MVAR simulation.

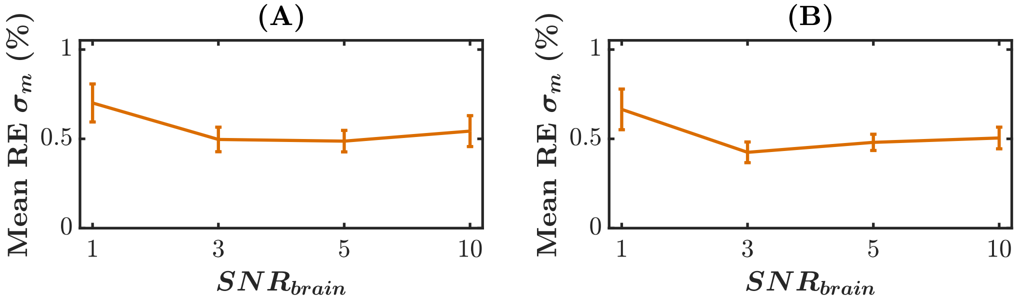


**Figure 7.** MVAR simulations. Mean reconstruction error of measurement noise ($\sigma_m$) for non-interacting (A) and unidirectionally interacting sources (B) for a range of $\mathrm{SNR}_{\mathrm{brain}}$. The error bars represent standard deviation, and the measurement SNR was fixed at 5.


3. F. Lindsten, P. Bunch, S. Särkkä, T. B. Schön, and S. J. Godsill. Rao-blackwellized particle smoothers for conditionally linear gaussian models. *IEEE Journal of Selected Topics in Signal Processing*, 10(2):353–365, 2016.

4. F. Lindsten, M. I. Jordan, and T. B. Schön. Particle gibbs with ancestor sampling. *Journal of Machine Learning Research*, 15:2145–2184, 2014.

5. K. B. Petersen, M. S. Pedersen, et al. The matrix cookbook. *Technical University of Denmark*, 7(15):510, 2008.

6. S. Särkkä. *Bayesian filtering and smoothing*, volume 3. Cambridge University Press, 2013.

7. A. Svensson, T. B. Schön, and F. Lindsten. Identification of jump markov linear models using particle filters. In *53rd IEEE Conference on Decision and Control*, pages 6504–6509, 2014.

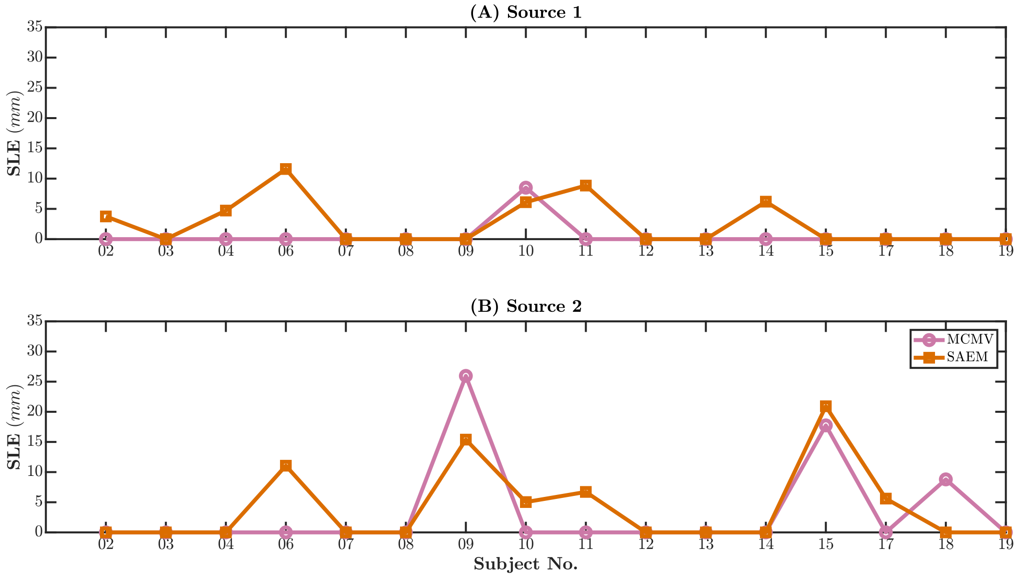


**Figure 8.** ECoG-based simulation. Source localization error in all subjects using the MCMV and SAEM approaches.

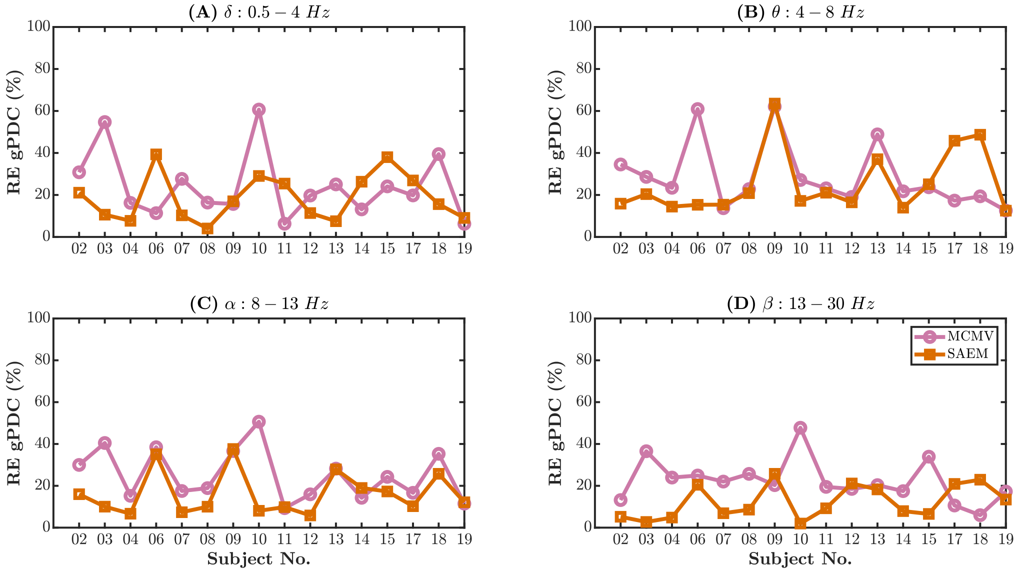


**Figure 9.** ECoG-based simulation. gPDC reconstruction errors in all subjects using the MCMV and SAEM approaches.
